# The role of toxin:antitoxin systems and insertion sequences in the loss of virulence in *Shigella sonnei*

**DOI:** 10.1101/2020.08.10.243006

**Authors:** Jessica E. Martyn, Giulia Pilla, Sarah Hollingshead, Mariya Lobanovska, Kristoffer S. Winther, Susan Lea, Gareth McVicker, Christoph M. Tang

## Abstract

The *Shigella* plasmid, pINV, contains a 30 kb pathogenicity island (PAI) encoding a Type III secretion system (T3SS) which is essential for virulence. During growth in the laboratory, avirulent colonies of *Shigella* (which do not express a T3SS) arise spontaneously. Avirulence in *Shigella flexneri* mostly follows loss of the PAI, following recombination between insertion sequences (ISs) on pINV; toxin:antitoxin (TA) systems on pINV promote its retention through post-segregational killing (PSK). We show that avirulence in *Shigella sonnei* mainly results from plasmid loss, consistent with previous findings; IS-mediated PAI deletions can occur in *S. sonnei*, but through different ISs than in *S. flexneri*. We investigated the molecular basis for frequent loss of the *S. sonnei* plasmid, pINV^*Ssonn*^. Introduction into pINV^*Ssonn*^ of CcdAB and GmvAT, toxin:antitoxin TA systems in pINV from *S. flexneri* but not *S. sonnei*, reduced plasmid loss and the emergence of avirulent bacteria. However, plasmid loss remained the leading cause of avirulence. We show that a single amino acid difference in the VapC toxin of the VapBC TA system in pINV also contributes to high frequency plasmid loss in *S. sonnei* compared to *S. flexneri*. Our findings demonstrate that the repertoire of ISs, complement of TA systems, and polymorphisms in TA systems influence plasmid dynamics and virulence loss in *S. sonnei*. Understanding the impact of polymorphisms should be informative about how TA systems contribute to PSK, and could be exploited for generating strains with stable plasmids.

## INTRODUCTION

*Shigella* spp. are important human pathogens as they are the leading cause of bacillary dysentery (Livio *et al.*, 2014). The Global Enteric Multicentre Study demonstrated that shigellosis is a major cause of moderate to severe diarrhoeal disease among children under five, with *Shigella flexneri* and *Shigella sonnei* accounting for around 90% of cases of shigellosis worldwide (Kotloff *et al.*, 2013, Livio *et al.*, 2014). *S. flexneri* is prevalent in lower:middle income countries, where individuals lack access to clean water and hygienic conditions (Kotloff *et al.*, 1999). In contrast, *S. sonnei* predominates in wealthier countries where the bacterium is largely transmitted from person-to-person (Kotloff *et al.*, 1999).

All species of *Shigella* have emerged from *Escherichia coli* following the acquisition of a non-conjugative ~ 210 kb plasmid, pINV (Lan & Reeves, 2002), which encodes a Type III secretion system (T3SS) on a 30 kb pathogenicity island (PAI) (Maurelli *et al.*, 1985, Sasakawa *et al.*, 1988). The T3SS delivers effector proteins, most of which are plasmid encoded, directly into host cells leading to bacterial invasion and immune evasion (Sansonetti *et al.*, 1986, Zychlinsky *et al.*, 1992, Schroeder & Hilbi, 2008, Mattock & Blocker, 2017). Furthermore, pINV from *S. sonnei* harbours genes responsible for the biosynthesis of the O antigen which is incorporated into the lipopolysaccharide (LPS) and capsule of the bacterium (Kopecko *et al.*, 1980, Sansonetti *et al.*, 1980, Xu *et al.*, 2002, Caboni *et al.*, 2015).

Seminal studies in the 1980s showed the contribution of pINV to *Shigella* virulence by characterising avirulent *S. sonnei* colonies that arise following loss of the plasmid during growth in the laboratory (Kopecko *et al.*, 1980, Sansonetti *et al.*, 1980, Sansonetti *et al.*, 1981). Interestingly, it was found that emergence of avirulence in *S. flexneri* is not associated with loss of the plasmid (Kopecko *et al.*, 1979). Instead, *S. flexneri* spontaneously loses expression of its T3SS following deletion of plasmid sequences encoding VirF, a key transcriptional regulator, and/or the entire T3SS PAI *via* intramolecular recombination between insertion sequences (ISs) (Schuch & Maurelli, 1997, Pilla *et al.*, 2017). ISs account for over 50% of open reading frames in *S. flexneri* pINV (pINV^*Sflex*^) (Buchrieser *et al.*, 2000, Venkatesan *et al.*, 2001). ISs also mediate the reversible integration of pINV^*Sflex*^ into the chromosome, providing the bacterium with bi-stable states of virulence/avirulence which mitigate the fitness costs of expressing the T3SS (Pilla *et al.*, 2017). Avirulent colonies of *S. flexneri* arise at a lower frequency than *S. sonnei*, with the mechanisms underlying the differences between the species largely unknown (Kopecko *et al.*, 1979, Kopecko *et al.*, 1980, McVicker & Tang, 2016).

pINV in *S. flexneri* harbours three functional toxin:antitoxin (TA) systems, VapBC (also known as MvpAT, for <u>m</u>aintenance of <u>v</u>irulence <u>p</u>lasmid AT), GmvAT, and CcdAB (Ogura & Hiraga, 1983, Radnedge *et al.*, 1997, Sayeed *et al.*, 2000, Sayeed *et al.*, 2005, McVicker & Tang, 2016, McVicker *et al.*, 2019). VapBC is the most well characterised TA system on pINV (Radnedge *et al.*, 1997, Sayeed *et al.*, 2000, Sayeed *et al.*, 2005, Dienemann *et al.*, 2011, McVicker & Tang, 2016, McVicker *et al.*, 2019); this TA system also contributes to antibiotic tolerance in *Escherichia coli* (Levin-Reisman *et al.*, 2017). The VapC toxin cleaves the initiator tRNA, tRNA^fMet^, preventing initiation of translation, while the VapB antitoxin binds VapC and blocks its activity (Dienemann *et al.*, 2011, Winther & Gerdes, 2011). VapBC is necessary and sufficient for retention of pINV^*Sflex*^ at 37°C (Sayeed *et al.*, 2005, McVicker & Tang, 2016), and also exerts local effects; repositioning *vapBC* next to the T3SS PAI reduces the occurrence of IS-mediated deletions of the T3SS PAI (Pilla *et al.*, 2017). GmvAT contributes to the maintenance of pINV^*Sflex*^ at environmental temperatures, while removal of CcdAB had no obvious effect on pINV under laboratory conditions; both GmvAT and CcdAB are absent from the *S. sonnei* virulence plasmid, pINV^*Ssonn*^, following ancestral deletions (McVicker & Tang, 2016).

Here we show that avirulent colonies in *S. sonnei* can result from discrete deletions involving the T3SS PAI as well as following plasmid loss. Deletions of the T3SS PAI in *S. sonnei* occur *via* different ISs than in *S. flexneri*, due to the distinct profiles of ISs in the plasmids from these species. However, in contrast to *S. flexneri,* loss of the T3SS PAI is not the most common cause of avirulence in *S. sonnei*. Consistent with previous work, loss of the plasmid is the major event that results in avirulent *S. sonnei* (Kopecko *et al.*, 1980, Sansonetti *et al.*, 1980, Sansonetti *et al.*, 1981). We show that pINV^*Ssonn*^ possesses two functional TA systems, RelBE and VapBC. However, RelBE has no discernible effect on pINV^*Ssonn*^ maintenance during growth in the laboratory. We also show that the high level of plasmid loss in *S. sonnei* was partially due to the absence of *ccdAB* and *gmvAT* from pINV^*Ssonn*^. Introduction of these *S. flexneri* TA systems into pINV^*Ssonn*^ significantly reduces the emergence of avirulence, but plasmid loss was still the most common molecular event leading to avirulence. We demonstrate that, in addition to the absence of CcdAB and GmvAT, a single amino acid substitution in VapC influences plasmid loss, with *S. flexneri* VapC stabilising pINV more efficiently than *S. sonnei* VapC through an unknown mechanism.

Plasmid loss poses a significant obstacle for studying *S. sonnei*. For example, a significant proportion of strains in genome sequencing studies lack pINV, hampering phylogenetic analyses of plasmid-encoded virulence factors (Holt *et al.*, 2012). Plasmid loss can also confound studies of host:pathogen interactions (Hartman & Venkatesan, 1998). Furthermore, live attenuated and whole cell vaccines are non-invasive and/or lack plasmid encoded antigens because of plasmid loss (Hartman & Venkatesan, 1998, Gerke *et al.*, 2015). Our findings highlight that the repertoire of ISs, the presence of TA systems, and an amino acid polymorphism in VapC have important consequences on plasmid dynamics in *S. sonnei*.

## RESULTS

### pINV^Ssonn^ has a RelBE homologue that does not contribute to plasmid stability in laboratory conditions

To define the mechanisms of pINV maintenance in *S. sonnei* 53G, we initially compared the sequence of its virulence plasmid with the plasmid from *S. flexneri* 5a M90T (Buchrieser *et al.*, 2000, Holt *et al.*, 2012). Whilst pINV^*Sflex*^ has three characterised TA systems, VapBC, GmvAT and CcdAB, pINV^*Ssonn*^ has ancestral deletions that result in the loss of *gmvAT* and *ccdAB* (McVicker & Tang, 2016). However, pINV^*Ssonn*^ has a predicted RelBE TA system, which is encoded near the origin of replication and is not present in pINV^*Sflex*^ (**Fig 1A**) (Jiang *et al.*, 2005). A RelBE TA system has been shown to stabilise a plasmid in *E. coli,* p307 (Gronlund & Gerdes, 1999). We examined genome sequences from a collection of 132 *S. sonnei* isolates obtained between 1948 and 2008 for the presence of *relBE*; only 46 of the 132 isolates have > 90 % sequence coverage of pINV due to plasmid loss (Holt et al., 2012). However, all 46 isolates with analysable plasmids contain *relBE* **(Table S1)**, demonstrating that this TA system is conserved in pINV^*Ssonn*^. Therefore, we further characterised pINV^*Ssonn*^ RelBE. *S. sonnei* RelE shares 87.5% amino acid identity with *E. coli* RelE, an mRNA-degrading toxin; RelB is 94.5% identical to *E. coli* RelB, which neutralises the toxicity of RelE (**Fig 1B**) (Gronlund & Gerdes, 1999). To determine whether pINV^*Ssonn*^ RelE is toxic, we expressed *relE* under the control of an arabinose-inducible promoter in *S. sonnei* 53G lacking pINV. Expression of *relE* alone led to a significant reduction in bacterial survival, consistent with RelE being toxic (*p* = 0.0028 comparing strains with and without *relE* induction, **Fig 1C**). Additionally, there was no difference in the *relBE* promoter from *E. coli* p307 and pINV^*Ssonn*^ **(Fig S1)** suggesting that the RelBE in *S. sonnei* is autoregulated by conditional cooperativity (Overgaard *et al.*, 2008). To assess whether RelB can rescue RelE-mediated toxicity, we induced *relE* expression in the presence or absence of *relB* expressed from its native promoter on a second plasmid, allowing modulation of RelB levels by conditional co-operativity (Overgaard *et al.*, 2008). The presence of *relE* under the control of its native promoter was sufficient to rescue toxicity following *relB* expression (**Fig 1C**), demonstrating that RelBE is a functional TA system.

**Figure 1.**
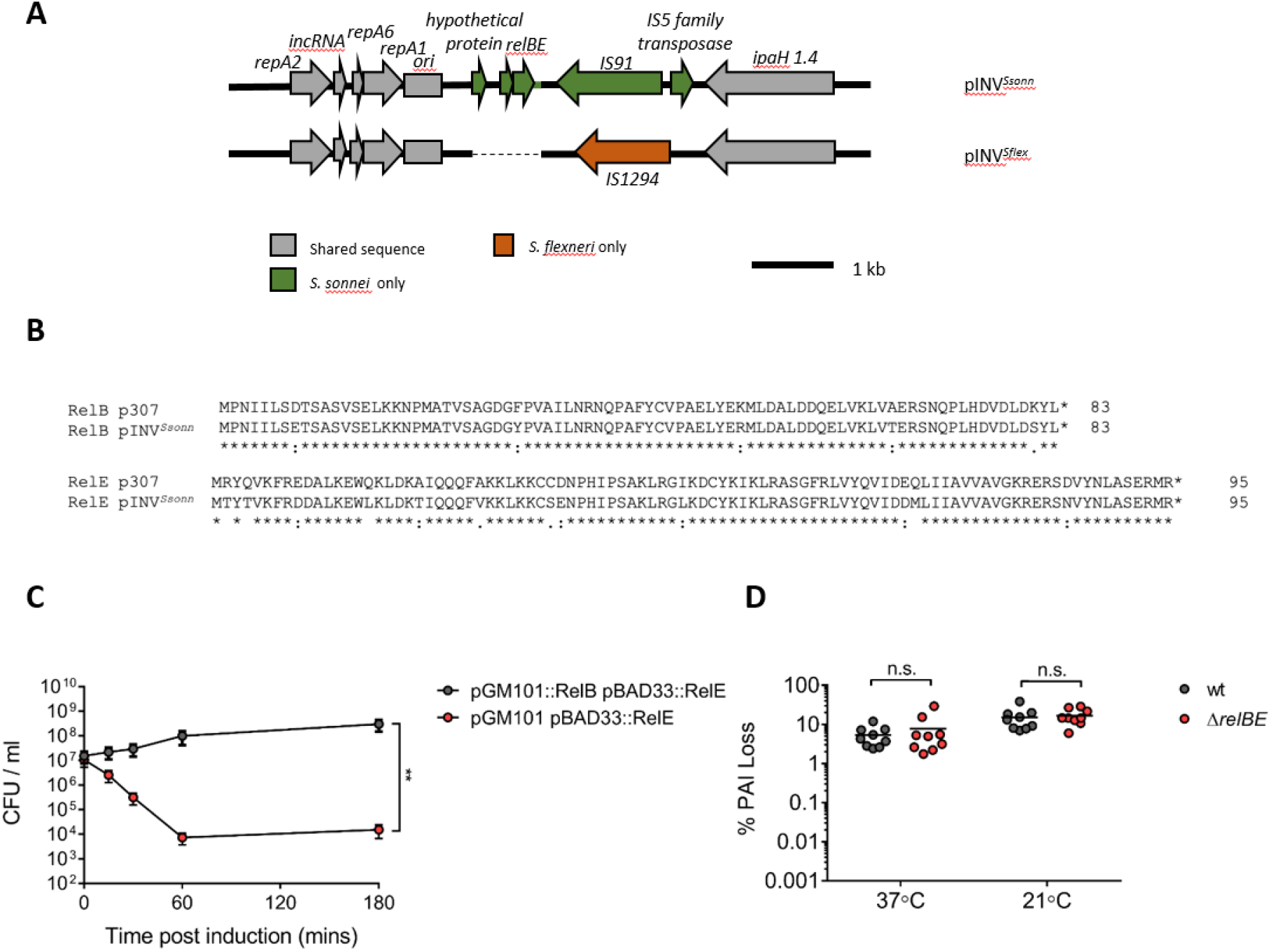
*S. sonnei* RelBE is a functional TA system that does not contribute to pINV^*Ssonn*^ stability. (A) Comparison of *S. flexneri* M90T pINV (AL391753) and *S. sonnei* 53G pINV (NC_016833) near the origin of replication (*ori*). (B) Alignment of RelBE from *E. coli* p307 (M26308) and from *S. sonnei* pINV (NC_016833) by Clustal O (1.2.4). Asterisks indicate identical amino acids, colons show amino acids with similar properties, and periods indicate amino acids sharing weakly similar properties. (C) Viability of *S. sonnei* 53G lacking pINV following expression of *relE* from pBAD33 in the presence/absence of *relB* on pGM101. **, *p* < 0.01 by two-way ANOVA with Sidak’s multiple comparisons test (n = 3 independent experiments, error bars show S.D.). (D) Loss of the PAI measured using *sacB*-*neo* on pINV^*Ssonn*^ with or without *relBE* following growth at 37°C or 21°C for ~ 25 generations. n.s. not significant by Mann-Whitney t-test (n = 9 colonies from three independent experiments).

To define the contribution of *relBE* to pINV^*Ssonn*^ maintenance, we deleted this TA system from pINV^*Ssonn*^ and measured plasmid stability using a *sacB-neo* assay as previously (McVicker & Tang, 2016). Briefly, we introduced a *sacB-neo* cassette into *mxiH* in the T3SS PAI of pINV^*Ssonn*^; after 25 generations of growth, bacteria that retain the T3SS PAI and harbour the *sacB-neo* cassette are resistant to kanamycin, while bacteria that have lost the marker (through deletions involving the T3SS PAI or loss of pINV) are resistant to sucrose (McVicker & Tang, 2016). Deletion of *relBE* had no significant impact on pINV maintenance during growth of *S. sonnei* at 37°C or 21°C (*p* = 0.9314 and 0.5891 respectively, **Fig 1D**), indicating that this TA system does not contribute to pINV^*Ssonn*^ maintenance under the conditions tested, although our results do not exclude the possibility that RelBE has a role *in vivo*.

### *Plasmid loss is the major event responsible for avirulence in* S. sonnei

Avirulent *Shigella* are distinguished from wild-type bacteria by the size and morphology of colonies growing on solid media containing the dye Congo red (CR) (Payne & Finkelstein, 1977, Maurelli *et al.*, 1984, Sasakawa *et al.*, 1986). Colonies of T3SS-expressing bacteria bind CR and appear red (CR^+^), while colonies of avirulent bacteria that have lost T3SS expression do not bind CR and therefore appear white (CR^−^). CR^−^ colonies are also larger than CR^+^ colonies, indicating that expression of the T3SS exerts a considerable metabolic cost (Schuch & Maurelli, 1997). We showed previously using a CR binding assay that a higher proportion of *S. sonnei* colonies are composed of avirulent bacteria than *S. flexneri* after 25 generations of growth (~ 5 % *vs*. ~ 0.05 %, respectively) (McVicker & Tang, 2016). Using assays with plasmids harbouring a *sacB-neo* cassette, we showed previously that introduction of *gmvAT* and *ccdAB* into pINV^*Ssonn*^ reduced the emergence of avirulent bacteria to similar levels as *S. flexneri* (McVicker & Tang, 2016). Although this assay is more sensitive at detecting low frequency events than using CR binding, it does not detect all genetic changes that lead to loss of virulence (*e.g.* loss of *virF* alone) (Schuch & Maurelli, 1997). Therefore, to gain an unbiased understanding of the molecular events underlying the loss of virulence in *S. sonnei*, we performed CR binding assays using wild-type *S. sonnei* and a strain with *ccdAB* and *gmvAT* on the plasmid (*ccdAB*^+^/*gmvAT*^+^). To prevent pINV^*Ssonn*^ loss before the start of experiments, we introduced a chloramphenicol resistance cassette (*cat*) into the plasmid in wild-type *S. sonnei* and *ccdAB*^+^/*gmvAT*^+^. Bacteria were initially grown in the presence of chloramphenicol, then transferred to antibiotic-free media and allowed to replicate for ~ 25 generations at 37°C. After this time, the emergence of CR^−^ colonies was quantified by plating bacteria to media containing CR. Consistent with previous results (McVicker & Tang, 2016), we found that the introduction of *gmvAT* and *ccdAB* into pINV^*Ssonn*^ significantly reduced the emergence of avirulent CR^−^ colonies (*p* = 0.0003 for *S. sonnei* +/− *ccdAB*/*gmvAT*, **Fig 2A**).

**Figure 2.**
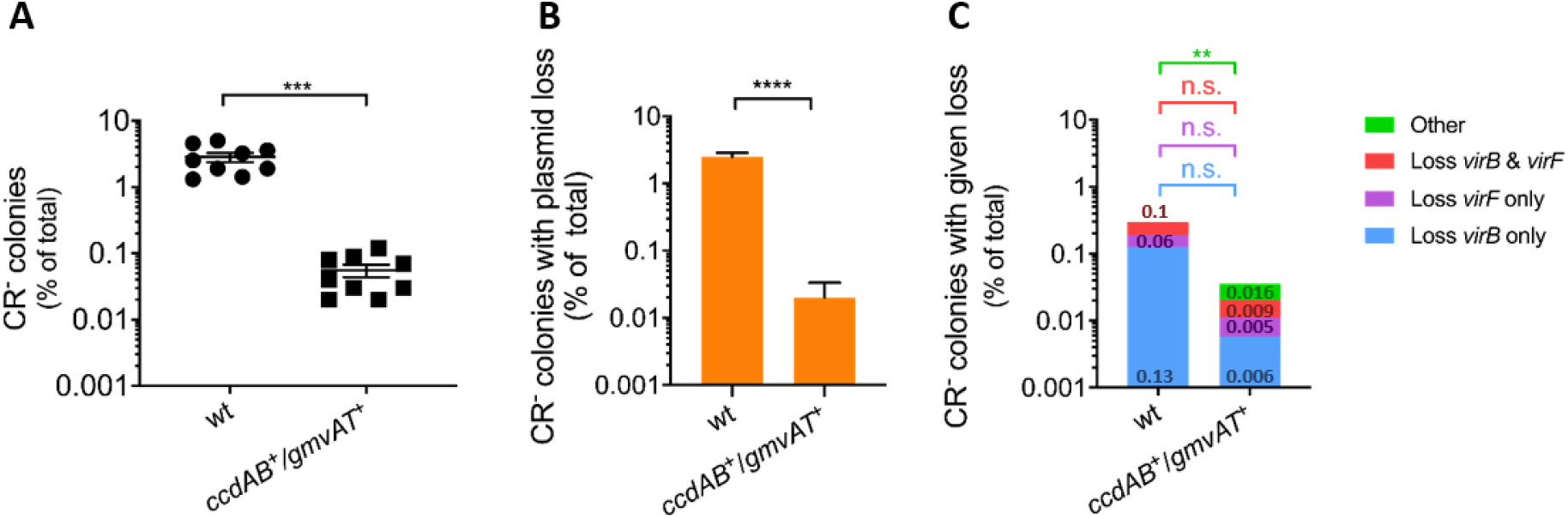
CR^−^ colonies of *S. sonnei* mostly result from loss of pINV. (A) The percentage of CR^−^ colonies emerging from wild-type *S. sonnei* 53G (wt) and a strain with *ccdAB* and *gmvAT* introduced into pINV (*ccdAB*^+^/*gmvAT*^+^) after growth at 37°C for ~ 25 generations. Solid line: mean (n = 9 biological replicates). ***, *p* ≤ 0.001 by parametric Welch’s *t*-test. (B and C) The number of CR^−^ colonies lacking specified virulence-related genes were assayed by multiplex PCR and shown as a % of all colonies. (B) Loss of the entire plasmid is inferred by loss of the origin of replication. (C) “Other” refers to CR^−^ colonies that contain *virB*, *virF* and the origin of replication. The value of each category is indicated inside the bars. Eight independent CR^−^ colonies were obtained from each of nine biological replicates (*i.e.* 72 colonies per strain). **, *p* ≤ 0.01 ****; *p* ≤ 0.0001 by non-parametric Mann-Whitney *t*-test. Error bars show S.D.

Next, we analysed CR^−^ bacteria by multiplex PCR to determine the molecular events leading to avirulence. We examined CR^−^ bacteria for the presence of the following elements: i) *virF*, which encodes a transcriptional regulator of T3SS PAI genes but is not in the T3SS PAI and; ii) *virB*, which encodes a transcriptional regulator located inside the PAI; iii) *ori*, the plasmid origin of replication which serves as a marker for pINV loss (Schuch & Maurelli, 1997, Pilla *et al.*, 2017); and iv) *hns*, which serves as a chromosomal control (**Fig 2B** and **C**). In contrast to *S. flexneri* (Pilla *et al.*, 2017), plasmid loss is the dominant cause of avirulence in wild-type *S. sonnei*, and is seen in approximately 2.5 % of all colonies after 25 generations (**Fig 2B**). However, similar to *S. flexneri* (Pilla *et al.*, 2017), deletion events affecting *virB* and/or *virF* were observed in avirulent bacteria emerging from wild-type *S. sonnei*, occurring in 0.3 % of cases (**Fig 2C**). The introduction of *ccdAB* and *gmvAT* into pINV^*Ssonn*^ significantly reduced plasmid loss, which dropped from 2.5 % to 0.02 % of all colonies in the absence/presence of *ccdAB* and *gmvAT,* respectively (*p* < 0.0001, **Fig 2B**). However, plasmid loss remained the main genetic event leading to avirulence in *ccdAB*^+^/*gmvAT*^+^, and was responsible for almost 50 % of avirulent colonies (**Fig 2B**). This demonstrates that although the introduction of *ccdAB* and *gmvAT*^+^ into pINV^*Ssonn*^ reduces the emergence of avirulent colonies, it is not sufficient to prevent plasmid loss.

### *ISs-mediated deletions of T3SS PAI-related genes on pINV*^Ssonn^ are distinct from those occurring on *pINV*^Sflex^

In *S. flexneri*, the most frequent molecular events leading to avirulence are deletions of the T3SS PAI with or without *virF* following intramolecular recombination between homologous copies of ISs (Schuch & Maurelli, 1997, Pilla *et al.*, 2017). To determine whether ISs also contribute to the loss of T3SS PAI-related genes from pINV^*Ssonn*^, we identified pairs of ISs in the same orientation and flanking the T3SS PAI and/or *virF*. PCR and sequence analysis revealed the involvement of three pairs of ISs in the loss of virulence: i) copies of IS*21*, (blue arc, **Fig 3A** - at positions 29,959–32,101 bp and 123,799–125,930 bp) mediated deletion of a 93.7 kb fragment, resulting in loss of the T3SS PAI and *virF*; ii) copies of IS*1* (grey arc, **Fig 3A** - at 73,733–74500 and 146,228–146996 bp) led to loss of the T3SS PAI; while iii) copies of IS*1294* (purple arc, **Fig 3A** at 33,932–35,621 and 68,896–70,585 bp) were involved in deletion of *virF* alone. Interestingly, IS*1* and IS*21* are not located near the T3SS PAI in pINV^*Sflex*^ (Buchrieser *et al.*, 2000) (**Fig 3B**), and were not found to mediate PAI deletions in *S. flexneri* (Pilla *et al.*, 2017). Although IS*1294* mediates loss of the T3SS PAI in pINV^*Sflex*^, loss of *virF* associated with this IS was not observed in *S. flexneri* previously (Pilla *et al.*, 2017). Therefore, ISs contribute to deletions affecting the T3SS PAI in *S. sonnei*, with different ISs involved compared with pINV^*Sflex*^.

**Figure 3.**
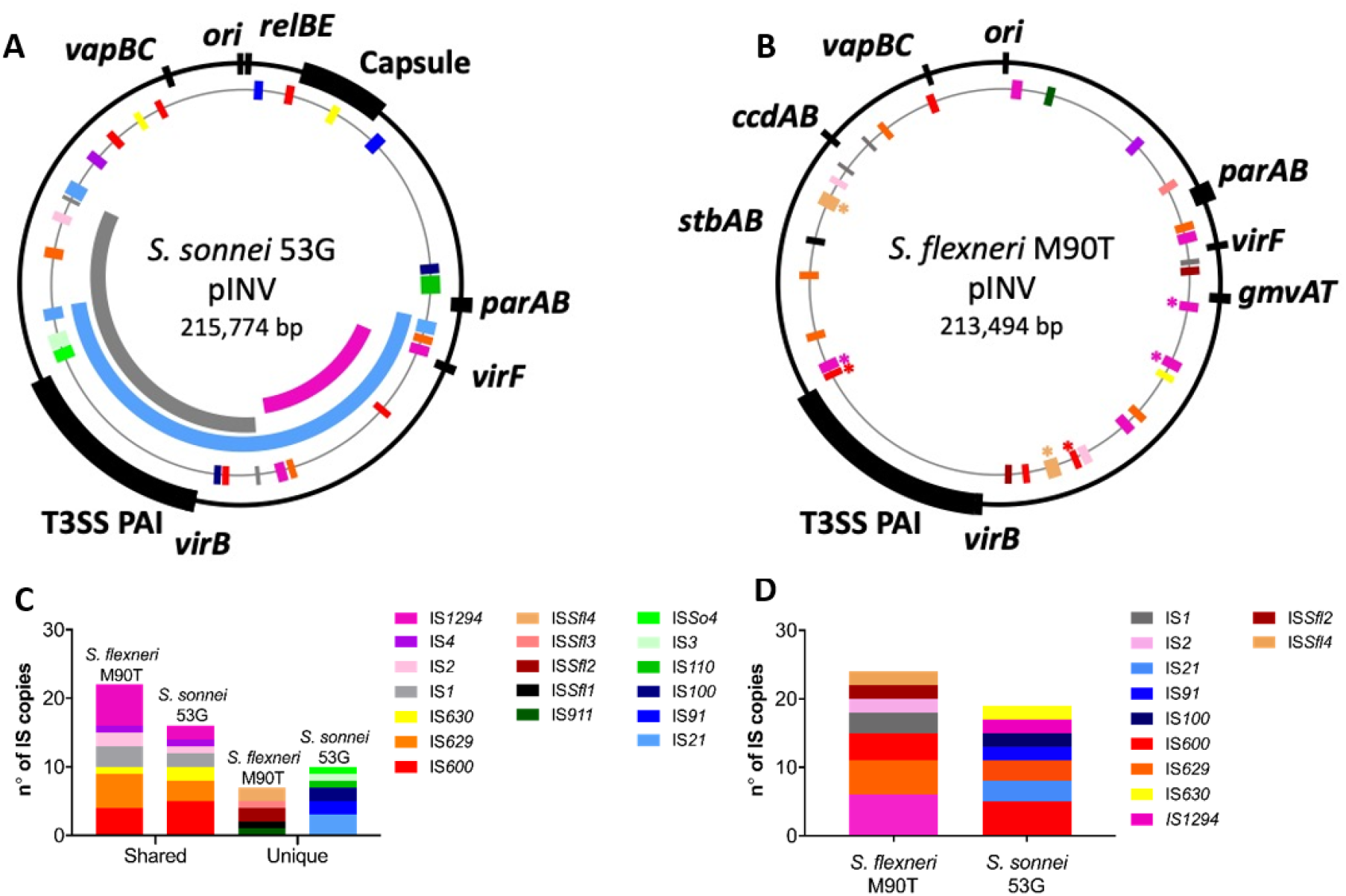
Distinct profile of ISs on pINV from *S. flexneri* M90T and *S. sonnei* 53G dictate different deletions of the T3SS PAI-related genes. (A) Alignment of three PAI deletions detected in *S. sonnei* 53G pINV (NC_016833, indicated as the black outer ring); insertion sequences are indicated as coloured boxes and follow the same colour-code used in panel B and C. (B) Schematic of the distribution of intact ISs on pINV from *S. flexneri* M90T (AL391753.1). ISs are coloured as indicated; asterisks indicate the ISs that mediate deletion of the T3SS PAI in *S. flexneri*. (C) Copies of ISs that are shared between the two strains, or only found in one strain (unique). (D) ISs that are present in more than one copy.

To examine whether differences in the deletions leading to avirulence in *S. flexneri* and *S. sonnei* are due to the presence of distinct ISs, we compared the content of ISs in pINV^*Sflex*^ and pINV^*Ssonn*^ (**Fig 3C and D**). In *S. flexneri* M90T pINV (pWR100), ISs account for 53 % of open reading frames, and IS*Sfl4,* IS*1294* and IS*600* are responsible for most T3SS PAI deletions (**Fig 3B**) (Buchrieser *et al.*, 2000, Pilla *et al.*, 2017). pINV^*Sflex*^ harbours a total of 29 different ISs, consisting of 12 ISs types. IS*1294* is the most abundant IS (present in six copies), with five copies of IS*629* and four of IS*600* on the plasmid, respectively; only four ISs are present in single copy (**Fig 3C** and **D**). In contrast, only ~ 23 % of open reading frames in pINV^*Ssonn*^ are intact ISs with 26 ISs belonging to 13 different types. IS*600* is the most abundant IS in pINV^*Ssonn*^, with five copies on the plasmid; the remaining ISs are present in up to three copies per plasmid, and seven are only present in single copy (**Fig 3C and D**). Therefore, there is a substantial difference in the repertoire and organisation of ISs on the two plasmids, with pINV^*Sflex*^ harbouring multiple copies of more ISs than pINV^*Ssonn*^.

### *A single amino acid substitution in VapC affects the rate of plasmid loss in* S. sonnei

We further investigated the reason for the residual plasmid loss following the introduction of *ccdAB* and *gmvAT* into pINV^*Ssonn*^. Given the importance of VapBC in the retention of pINV^*Sflex*^ (Radnedge *et al.*, 1997, Sayeed *et al.*, 2005, McVicker & Tang, 2016), we compared the nucleotide and predicted amino acid sequences of *vapBC* from *S. flexneri* M90T (AL391753) and *S. sonnei* 53G (NC_016833). We found that, while the promoter regions were identical, both VapB and VapC contain a single amino acid difference (**Fig S2**). The polymorphic residue in VapB (Thr^58^Ala, *S. flexneri vs. S. sonnei*) is located towards the C terminus of the antitoxin which is involved in neutralising VapC toxicity, while the polymorphic site in VapC (Lys^32^Arg, *S. flexneri vs. S. sonnei*) is not located near its active site or interface with VapB (Dienemann *et al.*, 2011). To determine if the *vapBC* sequences in *S. flexneri* M90T and *S. sonnei* 53G are representative of these species, we examined available *S. flexneri* and *S. sonnei* genome sequences for *vapBC* **(Tables S2 and S3)**. The predicted amino acid sequence of VapB in *S. flexneri* M90T is present in 95 % of sequenced *S. flexneri* strains, while VapB amino acid sequence in *S. sonnei* 53G is present in 87 % of strains **(Table S2).** Additionally, the VapC amino acid sequence of *S. flexneri* 5a M90T is present in 78 % of sequenced *S. flexneri* strains, and the amino acid sequence of *S. sonnei* 53G VapC is present in 82 % of strains **(Table S3).** Therefore, the predicted amino acid sequences of VapBC in *S. flexneri* M90T and *S. sonnei* 53G are representative of these species, and were designated VapBC^*Sflex*^ and VapBC^*Ssonn*^, respectively.

To define the effect of these substitutions on plasmid maintenance, we first used pSTAB as a model vector. pSTAB contains the origin of replication from pINV^*Sflex*^ along with a *sacB*-*neo* cassette that allows detection of the presence/absence of the plasmid without the confounding effect of other plasmid maintenance systems or ISs (McVicker *et al.*, 2019). We constructed derivatives of pSTAB containing *vapBC* from *S. sonnei* (generating pSTAB::VapBC^*Ssonn*^) or *S. flexneri* (pSTAB::VapBC^*Sflex*^), or containing chimeric *vapBC*s (*i.e.* pSTAB::VapB^*Sflex*^C^*Ssonn*^ or pSTAB::VapB^*Ssonn*^C^*Sflex*^). pSTAB derivatives were introduced into *S. sonnei* 53G lacking pINV, and plasmid loss was determined after ~ 25 generations. While pSTAB with *vapBC^Sflex^* was lost at a substantially lower rate compared with empty pSTAB (*p* < 0.0001, **Fig 4A**), introduction of *vapBC^Ssonn^* into pSTAB had no discernible impact on plasmid loss (*p* = 0.1873, pSTAB *vs.* pSTAB::VapBC^*Ssonn*^, **Fig 4A**). Furthermore, pSTAB containing *vapB^Ssonn^C^Sflex^* exhibited significantly reduced plasmid loss compared with empty pSTAB (*p* = 0.0002, pSTAB *vs.* pSTAB::VapB^*Ssonn*^C^*Sflex*^, **Fig 4A**). In contrast, *vapB^Sflex^vapC^Ssonn^* had no appreciable effect on pSTAB loss (*p* > 0.999, pSTAB *vs.* pSTAB::VapB^*Sflex*^C^*Ssonn*^, **Fig 4A**), indicating that the amino acid polymorphism in VapC (leading to a Lys^32^Arg substitution) affects the function of the VapBC TA system on plasmid maintenance in a model vector.

**Figure 4.**
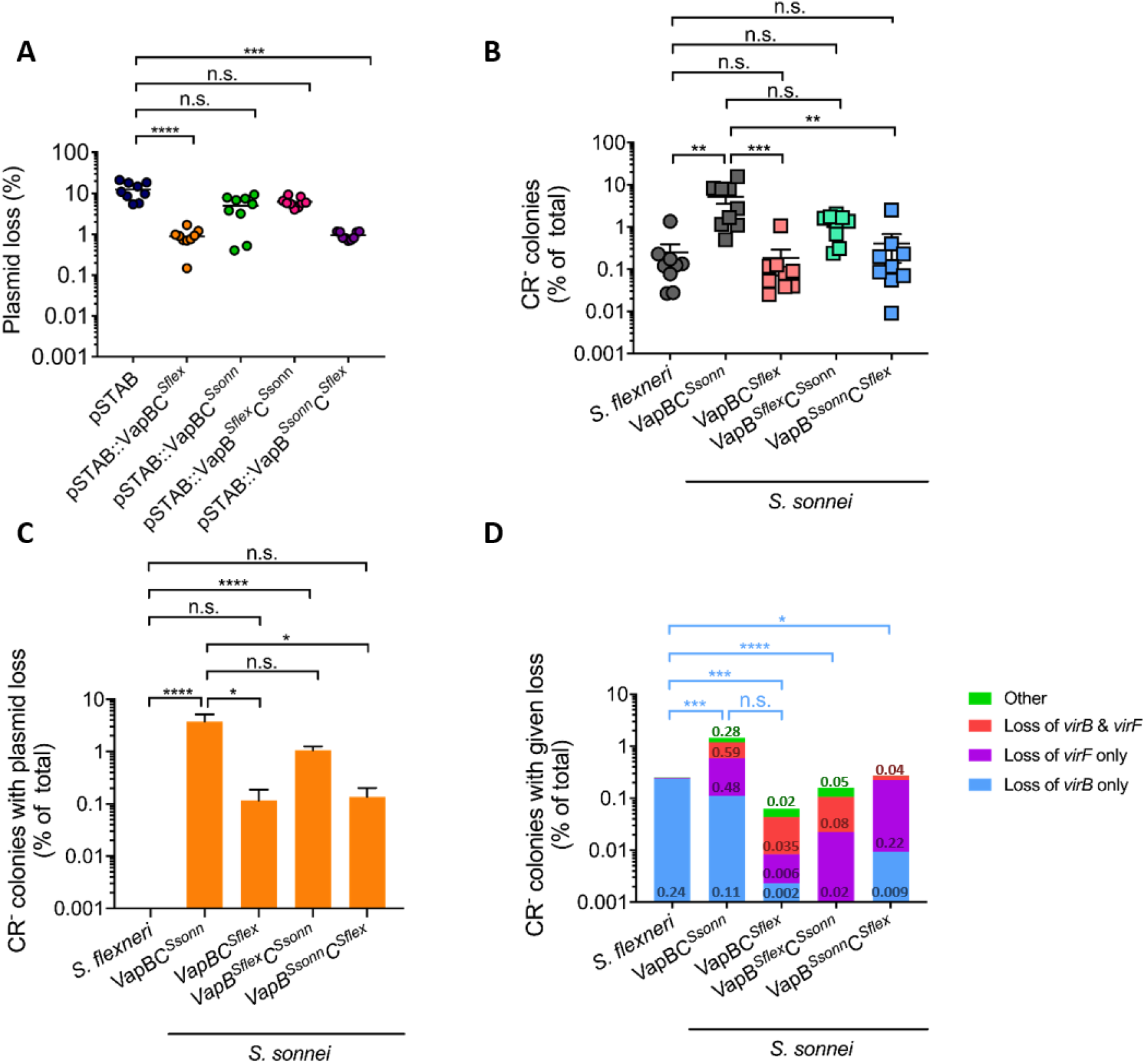
VapC polymorphisms affect plasmid maintenance. (A) The ability of VapBC^*Sflex*^, VapBC^*Ssonn*^, VapB^*Sflex*^C^*Ssonn*^, and VapB^*Ssonn*^C^*Sflex*^ to maintain pSTAB in *S. sonnei* 53G lacking pINV. Each dot represents the result for a single colony, and shows the percentage of all bacteria with plasmid loss. n = 9 for each strain which were tested on three independent occasions; horizontal lines show the mean. (B) The percentage of colonies that are CR^−^ of wild-type *S. flexneri* or *S. sonnei* (VapBC^Ssonn^), or *S. sonnei* with pINV^*Ssonn*^ harbouring *vapBC^Sflex^* (in strain VapBC^*Sflex*^), *vapB^Sflex^*C^*Ssonn*^ (in VapB^*Sflex*^C^*Ssonn*^) or *vapB^Ssonn^C^Sflex^* (VapB^*Ssonn*^C^*Sflex*^) after growth at 37°C for ~ 25 generations; solid line, mean + S.E.M. (n = 9 biological replicates). (C and D) The number of CR^−^ colonies lacking specified virulence-related genes were assayed by multiplex PCR and shown as a % of all colonies. (C) Loss of the entire plasmid is inferred by loss of the origin of replication; error bars, S.E.M. (D) “Other” refers to CR^−^ colonies that contain *virB*, *virF* and the origin of replication. The value of each category is indicated inside the bars. Eight independent CR^−^ colonies were obtained from each of nine biological replicates (*i.e.* 72 colonies were analysed per strain). ****, *p* ≤ 0.0001; ***, *p* ≤ 0.001; **, *p* ≤ 0.01; *, *p* ≤ 0.05; n.s. not significant; statistical comparison by one-way ANOVA with nonparametric analysis with Dunn’s multiple comparisons test.

Next, we investigated whether the VapC amino acid polymorphism also influences VapBC function in the context of pINV. We generated strains of *S. sonnei* containing pINV either with *S. sonnei vapBC* (generating strain VapBC^*Ssonn*^), *vapBC* from *S. flexneri* (strain VapBC^*Sflex*^), or a chimeric *vapBC* (*i.e.* VapB^*Sflex*^C^*Ssonn*^ and VapB^*Ssonn*^C^*Sflex*^); a chloramphenicol resistance cassette (*cat*) was inserted downstream of *vapBC* in all strains to ensure retention of pINV prior to the start of experiments. The emergence of CR^−^ colonies in these strains was quantified after growth for ~ 25 generations at 37°C in the absence of chloramphenicol. There was a significantly higher number of CR^−^ bacteria arising from VapBC^*Ssonn*^ compared with VapBC^*Sflex*^ or VapB^*Ssonn*^C^*Sflex*^ (*p* = 0.0003 and 0.0084, respectively, **Fig 4B**), similar to results with pSTAB. In contrast, there was no difference in the number of CR^−^ bacteria emerging from VapBC^*Ssonn*^ and VapB^*Sflex*^C^*Ssonn*^ (*p* > 0.9999, **Fig 4B**). We also investigated whether the single amino acid substitution in VapC led to a change in the molecular events that lead to avirulence. Of note, the presence of *vapC* from *S. flexneri* led to a significant reduction in plasmid loss as a cause of avirulence compared with *vapC* from *S. sonnei* (*p* = 0.010, VapBC*^Ssonn^ vs.* VapBC^*Sflex*^, and *p* = 0.0164, VapBC*^Ssonn^ vs.* VapB^*Ssonn*^C^*Sflex*^, **Fig 4C**). Additionally, there was a marked difference in the rate of plasmid loss in wild-type *S. flexneri* compared with wild-type *S. sonnei* or VapB^*Sflex*^C^*Ssonn*^ (*p* < 0.0001 for both comparisons, **Fig 4C**), but not when wild-type *S. flexneri* was compared with VapBC^*Sflex*^ or VapB^*Ssonn*^C^*Sflex*^ (*p* = 0.3882 and *p* = 0.2091, respectively, **Fig 4C**). Therefore, the presence of *S. flexneri vapC* leads to a significant decrease in the emergence of avirulence by reducing pINV loss, consistent with results obtained with pSTAB.

Furthermore, there were marked differences in the rates of *virB* and/or *virF* loss between wild-type *S. flexneri* and all *S. sonnei* strains. Loss of *virB* predominated in wild-type *S. flexneri* as previously (Pilla *et al.*, 2017), but was infrequently detected in *S. sonnei* strains (*p* = 0.0004, *S. flexneri vs*. VapBC^*Ssonn*^; *p* = 0.0006, *S. flexneri vs*. VapBC^*Sflex*^; *p* < 0.0001, *S. flexneri vs*. VapB^*Sflex*^C^*Ssonn*^, *p* = 0.0178, *S. flexneri vs.* VapB^*Ssonn*^C^*Sflex*^, **Fig 4D**). These findings provide further evidence that distinct IS-mediated deletions of the T3SS PAI-related genes occur in the plasmids from these two species because of their different IS profiles rather than any difference in VapBC.

As the genetic background can influence TA function (Cintron *et al.*, 2019), we also examined the effect of the VapC^*Sflex*^ polymorphisms in *S. flexneri*. We used tri-parental mating to transfer derivatives of pINV^*Ssonn*^ into *S. flexneri* BS176, which lacks pINV. The results of CR binding assays with pINV^*Ssonn*^ derivatives in *S. flexneri* demonstrate that the presence of *vapC^Sflex^* significantly reduces the emergence of CR^−^ colonies compared with *vapC^Ssonn^* (*p* = 0.0005, pINV^*Ssonn*^ VapBC*^Ssonn^ vs.* pINV^*Ssonn*^ VapBC^*Sflex*^, **Fig 5A**). Furthermore, multiplex PCR revealed that loss of the plasmid was not detected among 72 CR^−^ colonies analysed when *vapBC^Sflex^* was present on pINV^*Ssonn*^ in *S. flexneri* (*p* < 0.0001, for plasmid loss in *S. flexneri* with pINV^*Ssonn*^ VapBC*^Ssonn^ vs.* pINV^*Ssonn*^ VapBC^*Sflex*^, **Fig 5B**), with no change in the rate of *virB* and/or *virF* deletion (**Fig 5C**).

**Figure 5.**
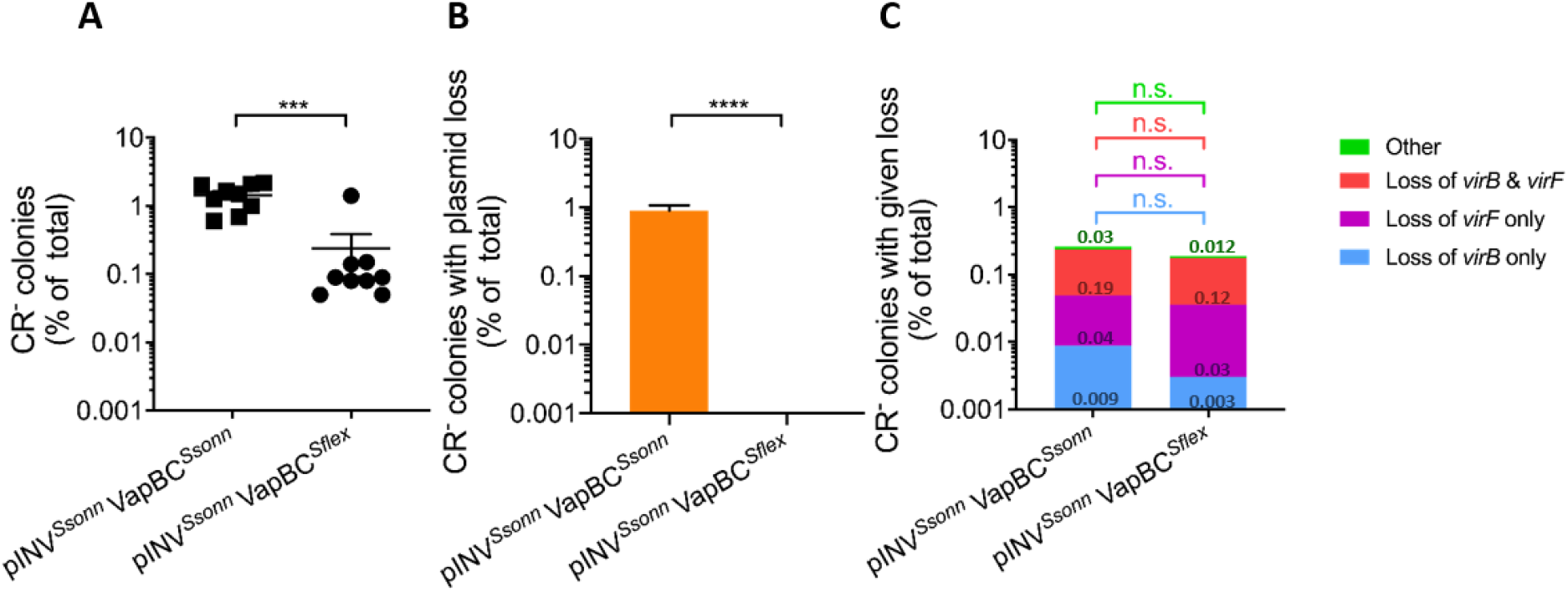
Analysis of *S. flexneri* carrying pINV ^*Ssonn*^ VapBC^*Ssonn*^ or pINV^*Ssonn*^ VapBC^*Sflex*^. (A) The percentage of colonies of *S. flexneri* carrying pINV^*Ssonn*^ VapBC^*Ssonn*^ or pINV^*Ssonn*^ VapBC^*Sflex*^ that are CR^−^ after growth at 37°C for approximately 25 generations; solid line, mean + S.E.M. (n = 9 biological replicates). (B and C) The number of CR^−^ colonies lacking lacking specified virulence-related genes detected by multiplex PCR and shown as a % of all colonies; error bars, S.E.M. (B) Loss of the entire plasmid is inferred by loss of the origin of replication. (C) “Other” refers to CR^−^ colonies that contain *virB*, *virF* and the origin of replication. The value of each category is indicated inside the corresponding region of the bars. Eight independent CR^−^ colonies were obtained from each of nine biological replicates (*i.e.* 72 colonies were analysed per strain). ****, *p* ≤ 0.0001; ***, *p* ≤ 0.001; **, *p* ≤ 0.01 *; *p* ≤ 0.05; n.s. not significant; statistical comparison by Mann-Whitney t-test.

Taken together, these results demonstrate that the single Lys^32^Arg substitution in VapC is responsible for the different ability of VapBC^*Ssonn*^ and VapBC^*Sflex*^ to prevent loss of pINV.

### Investigation of the effect of the VapC Lys^32^Arg substitution on VapBC structure:function

Next, we investigated the mechanism(s) by which the Lys^32^Arg substitution in VapC affects VapBC function. VapB and VapC act as proteins (Pullinger & Lax, 1992), and form a hetero-octomeric complex that binds its own promoter, leading to autoregulation (Robson *et al.*, 2009, Dienemann *et al.*, 2011, Winther & Gerdes, 2012). Therefore, we determined the atomic structure of VapBC from *S. sonnei* to establish whether the amino acid differences affect the overall architecture of the VapBC complex (**Fig 6A**, **Table 1**). Alignment of the *S. sonnei* and *S. flexneri* (PDB 3TND) VapBC hetero-octamers demonstrates that they are highly similar (root mean square deviation of 0.460 Å over all Cα atoms, **Fig 6A, Table 1**), indicating that the substitutions do not significantly alter the complex.

**Table 1.** Data collection and refinement statistics.

|  | VapBC (6SD6) |
| --- | --- |
| Wavelength |  |
| Resolution range | 35.15 - 2.61 (2.703 - 2.61) |
| Space group | P 32 2 1 |
| Unit cell | 95.8732 95.8732 116.389 90 90 120 |
| Total reflections | 38544 (3790) |
| Unique reflections | 19274 (1895) |
| Multiplicity | 2.0 (2.0) |
| Completeness (%) | 99.06 (97.32) |
| Mean I/sigma(I) | 9.82 (0.89) |
| Wilson B-factor | 71.32 |
| R-merge | 0.0358 (0.9819) |
| R-meas | 0.05062 (1.389) |
| R-pim | 0.0358 (0.9819) |
| CC1/2 | 0.999 (0.489) |
| CC* | 1 (0.811) |
| Reflections used in refinement | 19114 (1854) |
| Reflections used for R-free | 931 (85) |
| R-work | 0.2063 (0.3485) |
| R-free | 0.2359 (0.4159) |
| CC(work) | 0.964 (0.615) |
| CC(free) | 0.988 (0.559) |
| Number of non-hydrogen atoms | 3191 |
| macromolecules | 3164 |
| solvent | 27 |
| Protein residues | 402 |
| RMS(bonds) | 0.002 |
| RMS(angles) | 0.47 |
| Ramachandran favored (%) | 96.70 |
| Ramachandran allowed (%) | 3.30 |
| Ramachandran outliers (%) | 0.00 |
| Rotamer outliers (%) | 4.41 |
| Clashscore | 3.00 |
| Average B-factor | 80.58 |
| macromolecules | 80.65 |
| solvent | 72.81 |

Statistics for the highest-resolution shell are shown in parentheses.

**Figure 6.**
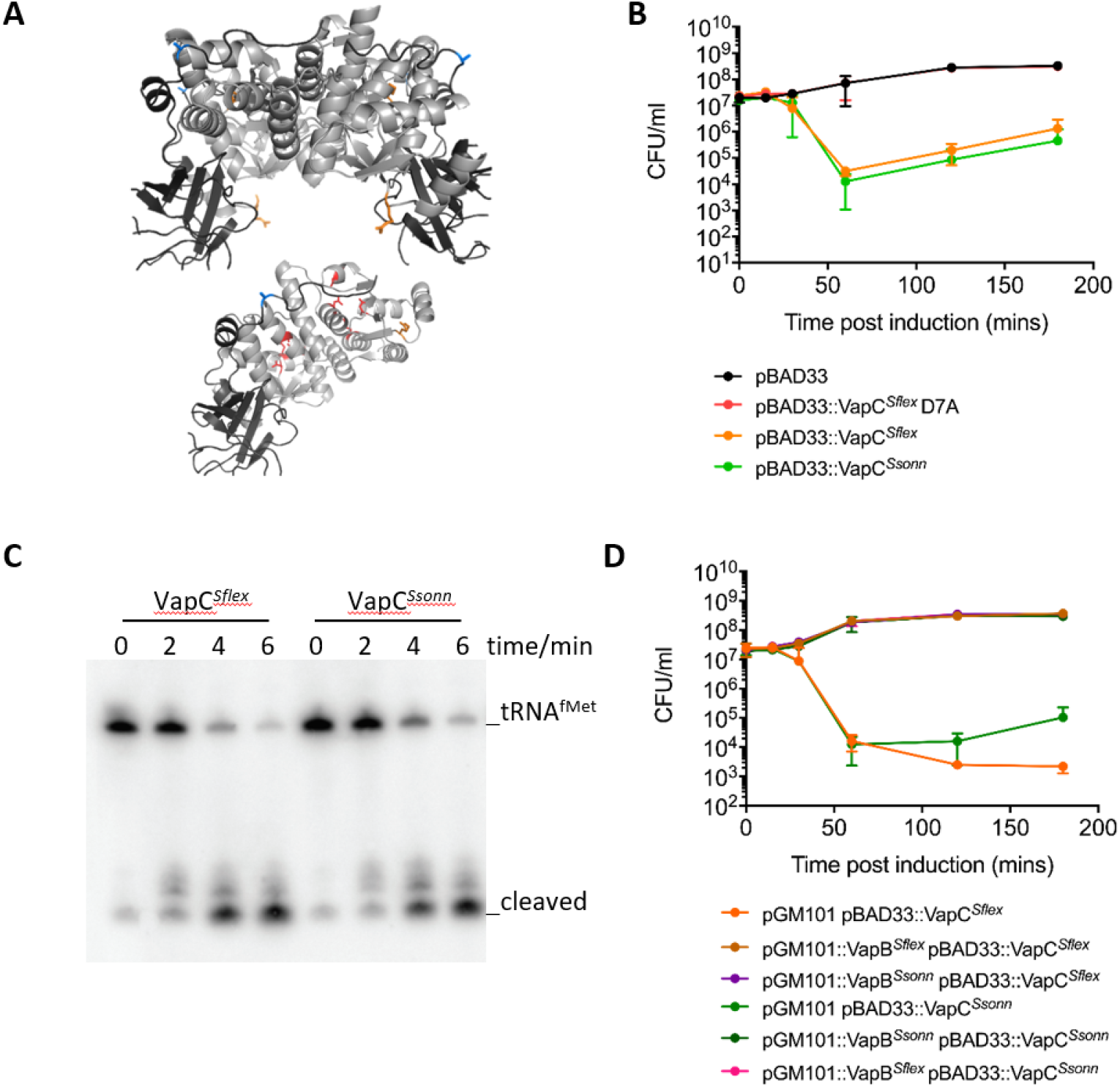
Analysis of the function of VapC polymorphisms. (A) Atomic structure of the *S. sonnei* 53G VapBC hetero-octomer (top) and the VapB^*Ssonn*^ dimer in complex with a VapC^*Ssonn*^ dimer (bottom); VapC^*Ssonn*^ is in light grey, and VapB^*Ssonn*^ is dark grey. The four residues comprising the VapC active site (D7, E42, D98 & E119) are highlighted in red. The location of the polymorphisms in VapB (T58A) and VapC (K32R) are highlighted in blue and orange, respectively. (B) Viability of *S. sonnei* 53G lacking pINV following expression VapC^*Ssonn*^ or VapC^*Sflex*^, or a non-functional version of VapC^*Sflex*^ (VapC^*Sflex*^D7A) from pBAD33; empty pBAD33, control. (C) Northern blot analysis of VapC-mediated cleavage of tRNA^fMet^ in *E. coli* MG1655 at times (indicated) following expression of VapC^*Ssonn*^ or VapC^*Sflex*^ from an inducible promoter. (D) Viability of *S. sonnei* 53G lacking pINV following expression of VapC^*Ssonn*^ or VapC^*Sflex*^ from pBAD33 with VapB^*Ssonn*^ or VapB^*Sflex*^ under the expression of their own promoters on pGM101. Error bars, S.D.

Next, we examined the effect of the Lys^32^Arg VapC substitution on its toxicity by expressing each version of VapC under the control of an arabinose-inducible promoter in *S. sonnei* lacking pINV. Induction of *vapC^Sflex^* (encoding VapC with Lys^32^) and *vapC^Ssonn^* (encoding VapC with Arg^32^) led to a similar reduction in bacterial survival from 1 hr post-induction onwards (*p* = 0.6867, pBAD33::VapC^*Ssonn*^ *vs.* pBAD33::VapC^*Sflex*^, **Fig 6B**). An inactive version of the toxin, VapC^*Sflex*^ D7A, was included as a negative control (**Fig 6B**) (Winther & Gerdes, 2011). We also assessed VapC activity by measuring cleavage of its target, tRNA^fMet^, using an *in vivo* cleavage assay as previously (Winther & Gerdes, 2011). VapC from *S. sonnei* or *S. flexneri* was expressed under an arabinose inducible promoter in *E. coli* MG1655, and the cleavage of tRNA^fMet^ was followed by Northern blot analysis. There was no detectable difference in the cleavage of tRNA^fMet^ by VapC^*Ssonn*^ and VapC^*Sflex*^ (**Fig 6C**).

Finally, we determined whether the VapC Lys^32^Arg substitution alters the capacity of VapB to act as its antitoxin. VapC was expressed under the control of an arabinose-inducible promoter in *S. sonnei* lacking pINV with either homologous or heterologous *vapB* under the control of its native promoter. There was no significant difference in the ability of VapB to neutralise VapC regardless of their source (*p* > 0.9999, **Fig 6D**).

## DISCUSSION

Here, we investigated the molecular mechanisms of pINV maintenance in *S. sonnei* and contrasted them with those in *S. flexneri* which does not lose its plasmid as frequently (McVicker & Tang, 2016). Three functional TA systems (VapBC, GmvAT and CcdAB) are distributed across *S. flexneri* pINV (Radnedge *et al.*, 1997, Sayeed *et al.*, 2000, Sayeed *et al.*, 2005, McVicker & Tang, 2016). In comparison *S. sonnei* has two TA systems (VapBC and RelBE), both of which are close to the origin of replication; VapBC is essential for pINV^*Sflex*^ retention at 37°C (Sayeed *et al.*, 2005, McVicker & Tang, 2016), and RelBE able to maintain plasmids in *E. coli* (Gotfredsen & Gerdes, 1998, Gronlund & Gerdes, 1999). Under the conditions tested, we were unable to detect any contribution of RelBE to the maintenance of pINV^*Ssonn*^ (**Fig 1D**). However, this does not exclude a role for this TA system in more physiologically relevant settings.

ISs are critical for the architecture and evolution of plasmids in *Shigella* (Buchrieser *et al.*, 2000, Venkatesan *et al.*, 2001). Previously, we observed that the spontaneous emergence of avirulence in *S. flexneri* is largely mediated by ISs and not plasmid loss (Pilla *et al.*, 2017). ISs can act as substrates for intramolecular recombination events, leading to gain/loss of genetic information. Indeed, recombination between ISs is likely to have resulted in the acquisition of the T3SS PAI (Venkatesan *et al.*, 2001), and causes deletions of PAI-associated genes in *S. flexneri* (Pilla *et al.*, 2017). We found that some avirulent *S. sonnei* also arise following deletions of PAI-associated genes but involve different ISs than *S. flexneri* (**Fig 3A**). Therefore, IS-mediated molecular rearrangements occur in *S. sonnei* leading to deletion of T3SS PAI-related genes, but these events often go undetected because of the high level of plasmid loss (**Fig 2B**). Differences in ISs in pINV^*Ssonn*^ and pINV^*Sflex*^ (**Fig 3**) might also affect the repertoire of T3SS effectors in the two species.

Previously, we found that ancestral deletions of two TA systems, GmvAT and CcdAB, decrease the stability of pINV^*Ssonn*^ (McVicker & Tang, 2016). However even when GmvAT and CcdAB were introduced into pINV^*Ssonn*^, ~ 50 % of avirulent bacteria still resulted from plasmid loss (**Fig 2B**). VapBC (also referred to as MvpAT) is the only TA system present in pINV from both *S. flexneri* and *S. sonnei*, and plays a critical role in maintaining pINV^*Sflex*^ (Radnedge *et al.*, 1997, Sayeed *et al.*, 2000, Sayeed *et al.*, 2005, McVicker & Tang, 2016). Given this, we hypothesised that differences in VapBC could also contribute to the high plasmid loss in *S. sonnei*. We found that a single amino acid polymorphism in VapC has a profound effect on the function of this TA system in pSTAB and pINV. The predicted sequences of VapB and VapC in *S. flexneri* M90T and *S. sonnei* 53G are found in most sequenced isolates (**Table S2** and **S3**), so our results are broadly applicable to these two species.

Even though VapBC^*Ssonn*^ is less efficient at post-segregational killing (PSK) than VapBC^*Sflex*^ (**Fig 4 and 5**), we could not delete this TA system from pINV^*Ssonn*^ despite multiple attempts. It is likely that VapBC is essential for retention of pINV^*Ssonn*^ (McVicker & Tang, 2016).

The amino acid substitution in VapC could affect plasmid loss by several mechanisms during PSK. Of note, the amino acid change in VapC is not in its PIN (PilT N-terminus) domain, which is required for tRNA^fMet^ cleavage (Winther & Gerdes, 2011), or in the region that interacts with VapB (**Fig 6A**) (Dienemann *et al.*, 2011). To investigate how the VapC polymorphism influences plasmid loss, we determined the atomic structure of the *S. sonnei* VapBC complex. However, the amino acid substitutions do not affect the overall structure of VapBC. Furthermore, there was no detectable change in the toxicity of VapC, its ability to cleave tRNA^fMet^, or its susceptibility to be neutralised by VapB (**Fig 6B, C and D**). It is possible that these assays are not sufficiently sensitive to detect subtle yet important differences in VapC activity and/or its interaction with VapB. Further studies are underway to identify other potential changes in VapC function associated with the amino acid substitution. Also, it is possible that other sequence variants of VapBC (**Table S2 and S3**) alter the activity of this TA system.

TA systems can stabilise local sequences by preventing IS-mediated deletions (Pilla *et al.*, 2017). Our results suggest that GmvAT might reduce IS*1294*-mediated loss of *virF* in *S. flexneri*, as this system is located only 7.4 kb from *virF* on pINV^*Sflex*^ within a region enclosed by two copies of IS*1294* (Buchrieser et al., 2000). The absence of *gmvAT* from pINV^*Ssonn*^ might explain our finding that avirulent strains lacking only *virF* are more common in *S. sonnei* than *S. flexneri* (**Fig 2C**). This hypothesis is supported by the decrease in *virF* loss seen in pINV^*Ssonn*^ containing *gmvAT* (**Fig 2C**).

*S. sonnei* is a highly successful pathogen despite having an unstable virulence plasmid (Kopecko *et al.*, 1980, Sansonetti *et al.*, 1980, Sansonetti *et al.*, 1981). The emergence of avirulent bacteria generates heterogeneity, which might be beneficial for the survival of *S. sonnei* in certain circumstances. For example, rapidly replicating avirulent bacteria could act as an immune decoy in the intestinal tract, or provoke local inflammatory responses to modify the local micro-environment in favour of their virulent relatives (Tsai & Coombes, 2019). Additionally, IS-mediated deletions of the T3SS PAI could be a trade-off for genetic plasticity that allows the acquisition of novel traits, such as genes encoding T3SS effectors, with ISs acting as substrates for recombination during horizontal gene transfer.

Understanding differences between pINV in *S. sonnei* and *S. flexneri*, and the influence of TA systems, such as GmvAT, CcdAB and VapBC, and ISs could explain the epidemiology of these two important pathogens. Also, further studies of VapBC should elucidate how the Lys^32^Arg substitution in VapC affect the function of this important TA system. Our findings could be exploited to construct strains of *S. sonnei* with more stable plasmids. This would facilitate molecular studies of this bacterium and may aid the development of vaccines and animal/human challenge models. Vaccines based on live attenuated *S. sonnei* strains can be adversely influenced by plasmid loss (Hartman & Venkatesan, 1998, Kotloff *et al.*, 2002, Barnoy *et al.*, 2011, Bedford *et al.*, 2011). Plasmid loss can also affect other vaccine strategies, including those based on outer membrane vesicles, by reducing the amount of immunogenic plasmid-encoded T3SS effectors (Gerke *et al.*, 2015), while detailed knowledge of plasmid dynamics might offer novel approaches to target the plasmid maintenance systems to combat this important human pathogen. Finally, understanding the impact of polymorphisms in TA systems should be informative about the process of PSK, and persistence/tolerance which can also be mediated by TA systems.

## EXPERIMENTAL PROCEDURES

### Bacterial strains and growth

Bacterial strains and plasmids used in this study are shown in **Tables S4** and **S5**, respectively. *Shigella* spp. and *E. coli* were grown in TSB (Tryptic soy broth, Sigma) and LB (Lysogeny broth, Invitrogen), respectively; 1.5% (wt./vol.) agar (Oxoid) was added for solid media. Antibiotics were used at the following concentrations: carbenicillin, 100 μg/ml; chloramphenicol, 5 μg/ml (for *S. flexneri*) and 20 μg/ml (for *S. sonnei*); kanamycin, 50 μg/ml; streptomycin, 100 μg/ml. Congo red (CR, 0.01% wt./vol., final concentration) was added to TSB agar (CR-TSA). For sucrose selection, 1% (w/v) bacto-tryptone (Sigma), 0.5% (wt./vol.), yeast extract (Sigma) and agar were autoclaved in water, and sucrose (VWR, final concentration of 10%, wt./vol.) added prior to pouring plates.

### Construction of strains

Lambda Red recombination was employed to construct mutants (Datsenko & Wanner, 2000). Approximately 1 kb of sequence upstream and downstream of the gene interest was used to flank an antibiotic resistance cassette. Fragments were amplified by PCR (primers used in this study are shown in **Table S6**) then ligated into pUC19 (Yanisch-Perron *et al.*, 1985) using NEBuilder^®^ HiFi master mix (New England Biolabs). Resulting plasmids were transformed into *E. coli* DH5α, and used as the template to generate approximately 1 μg of linear DNA by PCR, which was transformed by electroporation into *S. sonnei* 53G expressing the recombinase from pKD46 (Datsenko & Wanner, 2000). Bacteria were plated onto CR-TSA containing appropriate antibiotics and incubated overnight at 37°C. Strains were checked by PCR and sequencing.

For construction of pET28a-VapBC^*Ssonn*^, *vapB* and *vapC* were amplified from *S. sonnei* genomic DNA using the primers described in **Table S6**, then inserted into pET28a digested with *Nde*I and *Xho*I using NEBuilder^®^ HiFi master mix.

### Tri-parental mating

To transfer pINV^*Ssonn*^ into *S. flexneri*, tri-parental mating was performed using *S. sonnei* 53G with a *cat* cassette downstream of *vapBC* (conferring chloramphenicol resistance) as the donor strain, *S. flexneri* BS176 (streptomycin resistant) as the recipient, and *E. coli* containing pRK2013 (conferring kanamycin resistance) as the helper strain (Figurski & Helinski, 1979, Ditta *et al.*, 1980). Strains were grown separately overnight at 37°C in 10 ml LB liquid medium with appropriate antibiotics. Cultures were re-suspended in 10 ml PBS after washing, then mixed at a ratio of 1:1:5 of donor, helper, and recipient, respectively, in a 100 μl volume. Bacteria were spotted onto LB agar without antibiotics and incubated overnight at 37°C. Bacteria were harvested from spots and re-suspended in 1 ml TSB, and finally plated onto CR-TSA containing chloramphenicol and streptomycin to select for transconjugants.

### sacB-neo *assays*

*S. sonnei* strains possessing pINV with the *sacB*-*neo* cassette in *mxiH* or pSTAB derivatives were grown from frozen stocks at 37°C for 16 hours or at 21°C for 96 hours when plated on solid LB medium to reach ~ 25 generations. On three separate occasions, three colonies were resuspended in 100 μl PBS and serial dilutions plated on solid media with sucrose or with kanamycin only. PAI loss was calculated as the ratio of CFU on plates with sucrose and on plates without sucrose/with kanamycin, and shown as a percentage.

### CR-binding assays

*Shigella* spp. were grown at 37°C on CR-TSA plates containing chloramphenicol overnight to obtain single colonies. On three separate occasions, three independent CR^+^ colonies were re-suspended in 5 ml TSB liquid medium and incubated at 37°C with shaking at 180 r.p.m. for 16 hr (~ 25 generations). Samples were diluted in PBS and plated onto CR-TSA, and incubated overnight at 37°C before counting the number of CR^+^ and CR^−^ colonies. The proportion of CR^−^ colonies was quantified by dividing the number of emerging CR^−^ colonies by the total number of colonies (CR^+^ and CR^−^), and expressed as a percentage.

### Multiplex PCR

Multiplex PCR was used to detect *virF*, *virB*, and *ori* (the pINV origin of replication); *hns* was included as a positive chromosomal control. Reactions included Taq polymerase (Sigma Aldrich) with an annealing temperature of 51.2°C and extension time of 1.5 min. For each strain, eight CR^−^ colonies emerged from each biological repeat following the CR binding assay were analysed. For each strain, the % of all colonies showing a given gene loss was calculated by first quantifying the % of CR^−^ colonies with a particular gene loss and then adjusting this result using the % of CR^−^ colonies in the total bacterial population as described in the CR binding assay.

### Toxicity, anti-toxicity and VapC-mediated cleavage assays

To assess the toxicity of VapC, the protein was expressed using pBAD33 (Guzman *et al.*, 1995) in *S. sonnei* 53G lacking pINV. Initially, bacteria were grown at 37°C with shaking at 180 r.p.m in LB liquid medium with 0.2 % glucose (wt./vol.) to repress toxin expression. At an optical density (OD) A_600_ ~ 0.1 cultures were pelleted by centrifugation at 3000 *x g* then re-suspended in LB liquid medium with 0.2% arabinose (wt./vol.) to induce toxin expression, and grown at 37°C with shaking at 180 r.p.m.. Aliquots of cultures were taken at time points following induction then serially diluted in PBS and plated onto LB solid media containing 0.2 % glucose (wt./vol.) to measure bacterial viability.

To assess the ability of VapB to prevent VapC toxicity, VapB from *S. flexneri* or *S. sonnei* was expressed from their native promoter on pGM101 in *S. sonnei* 53G lacking pINV with or without pBAD33::*vapC^Sflex^* or pBAD33::*vapC^Ssonn^*(McVicker & Tang, 2016). Expression of VapC was induced as above in bacteria with or without a plasmid containing *vapB*, and viability was assessed by plating aliquots of cultures to solid media with 0.2 % glucose (wt./vol.).

To assess the ability of VapC to cleave tRNA^fMet^, *E. coli* MG1655 containing pBAD33::VapC^*Sflex*^ or pBAD33:: VapC^*Ssonn*^ were grown exponentially in LB liquid medium at 37°C with shaking at 180 r.p.m. At an O.D. A_600_ ~ 0.4, toxin expression was induced by addition of 0.2% arabinose (final concentration, wt./vol.). Samples (1 ml) were collected before (0 min) and 2, 4 and 6 min after addition of arabinose, and immediately mixed with 125 μl of 5 % phenol in ethanol on ice to prevent further RNA degradation. Samples were harvested by centrifugation at 3000 *x g* for 5 min at 4°C, and RNA extracted using the hot phenol method as previously (Winther *et al.*, 2013). Total RNA (2.5 μg) was denatured in formamide and separated on a denaturing 8% polyacrylamide gel (19:1) containing 8M urea buffered in Tris-Borate-EDTA (TBE); the RNA was transferred to a Zeta-Probe membrane (Bio-Rad) by electroblotting. Membranes were pre-hybridized in hybridization buffer (0.9M NaCl, 0.05M NaH_2_PO_4_, 0.05M EDTA, 5 x Denhardt’s solution (ThermoFischer Scientific), 0.5 % SDS and 550 μg salmon sperm DNA, pH 7.4) for 30 min at 42°C before addition of the DNA probe. The probe was generated by 5’ phosphorylation of 30 pmol of tRNA^fMet^ specific DNA oligonucleotide (5’-CTTCGGGTTATGAGCCCGACGAGCTA) with 30 μCi [^32^P]-ATP using 1 μl T4 polynucleotide kinase (ThermoFischer Scientific) in a total volume of 20 μl according to manufacturer’s instructions. Hybridisation was continued overnight at 42°C. To reduce non-specific hybridization, membranes were washed in 2 x SSC (0.3M NaCl, 0.03M Na_3_C_6_H_5_O_7_) with 0.1% SDS at room temperature. Cleavage of tRNA^fMet^ was visualised by phosphor imaging.

#### Protein purification and crystallography

pET28a-VapBC^*Ssonn*^ was transformed into *E. coli* C41. Following growth at 37°C to OD_600_ ~ 0.8, expression was induced with 1mM IPTG and cultures incubated for a further three hours prior to harvesting by centrifugation at 5000 *x g* for 10 minutes. VapBC was purified as described previously (Dienemann *et al.*, 2011). In brief, 150 mM VapBC^*Ssonn*^ octamer was combined with double stranded DNA (5’-ACAATAGATATACACAAGACATATCCAC-3’) re-suspended in H_2_O using a ratio of 1:1.2 of VapBC^*Ssonn*^ octamer to DNA. The VapBC^*Ssonn*^:DNA mixture was dialysed in a solution of 25 mM Tris pH 8.0 for five hrs at room temperature in a slide-alyser cassette with a 3.5 kDa cut-off. Crystals were grown using the sitting drop method in 0.1M ammonium sulphate, 0.3M sodium formate, 0.1M sodium cacodylate pH 6.5, 3% γ-PGA and 5% PEG 4000 at a ratio of 0.4:0.6 of protein to mother liquor. Crystals obtained were formed of VapBC^*Ssonn*^ alone. Crystals were briefly incubated in a solution of crystallisation buffer supplemented with 40 % ethylene glycol, followed by flash freezing in liquid nitrogen. Data were collected on beamline I04-1 at Diamond Light Source and indexed, scaled and reduced using x2 dials (Winter *et al.*, 2018) within ISPYB (Delageniere *et al.*, 2011). The structure was solved by molecular replacement using PHASER (McCoy *et al.*, 2007) within CCP4i (Winn *et al.*, 2011) with the structure of VapBC^*Sflex*^ (3TND, (Dienemann *et al.*, 2011)) as the starting model. Iterative manual rebuilding and refinement using Coot (Emsley *et al.*, 2010) and PHENIX (Adams *et al.*, 2010) led to the models described in **Table 1**. Coordinates and structure factors have been deposited in the Protein Data Bank (accession number, ID 6SD6).

### Bioinformatics

Schematics of the position of RelBE or IS pairs on pINV from *Shigella* spp. were created from the sequence of S. *flexneri* M90T pINV (pWR100, accession number AL391753.1), and *S. sonnei* 53G pINV (accession number, NC_016833). To determine the presence/absence of RelBE in a collection of *S. sonnei* from Holt *et al.* 2012 BLAST (BLASTn) v2.9.0 was used. Alignment of RelBE was performed using BLAST (BLASTn and BLASTp) v2.9.0 and subsequently Clustal O of sequence from RelBE from p307 from *E. coli* (accession number, M26308) and *S. sonnei* 53G pINV (accession number, NC_016833). Polymorphisms in VapB and VapC were identified in available *S. flexneri* or *S. sonnei* plasmid sequences using BLASTp (v2.9.0) and alignment with Clustal O (v1.2.4).

### Statistical analyses

All data were analysed using Graphpad Prism (version 7.4). Data were first tested for normal distribution using the Sharpiro Wilkes test. If the data were not normally distributed then a non-parametric one-way ANOVA was performed. If data were normally distributed then a Gaussian distribution was assumed, and a parametric one-way ANOVA was performed. Alternatively, a non-parametric t-test was performed using a Mann-Whitney test.

## ACKNOWLEDGEMENTS

We are grateful for the support from the Wellcome Trust (CMT), and the EP Abraham Trust and an MRC studentship (to GP). We thank Diamond Light Source for beamtime (proposal mx12346), and the staff of beamline I04-1 for assistance with crystal testing and data collection. The authors declare no conflicts of interest, and comply with the Expects Data Sharing Policy.

---

**Supplementary Figure 1 Sequence alignment of the RelBE promoter** Alignment of the promoter region of *relBE* Alignment of RelBE from p307 from *E. coli* (M26308) and pINV from *S. sonnei* (NC_016833) by CLUSTAL O (1.2.4). Asterisks indicate identical nucleotides. The predicted −10 and −35 sequences are highlighted in bold. The putative *relO* sites are indicated highlighted. The first codon of RelB is indicated using a M.

**Supplementary Figure 2 Polymorphisms in VapBC.** Alignment of the *vapBC* promoter (A) VapB (B) and VapC (C) from *S. flexneri* M90T pINV (AL391753) and *S. sonnei* 53G pINV (NC_016833) by CLUSTAL O (1.2.4). Polymorphic residues are shown in red. Asterisks indicate identical nucleotides/amino acids. The predicted −10 and −35 sequences are highlighted in bold. The putative *vapO* sites are indicated highlighted. The first codon of VapB is indicated using a M.

**Supplementary Table 1: The presence of *relBE* on pINV^*Ssonn*^**

**Supplementary Table 2: Polymorphisms in VapB from *S. flexneri* and *S. sonnei.***

**Supplementary Table 3: Polymorphisms in VapC from *S. flexneri* and *S. sonnei.***

**Supplementary Table 4: Bacterial strains used in this study**

**Supplementary Table 5: Plasmids used in this study**

**Supplementary Table 6: Primers used in this study**

## REFERENCES

Adams, P.D., Afonine, P.V., Bunkoczi, G., Chen, V.B., Davis, I.W., Echols, N., Headd, J.J., Hung, L.W., Kapral, G.J., Grosse-Kunstleve, R.W., McCoy, A.J., Moriarty, N.W., Oeffner, R., Read, R.J., Richardson, D.C., Richardson, J.S., Terwilliger, T.C., and Zwart, P.H. (2010) PHENIX: a comprehensive Python-based system for macromolecular structure solution. Acta Crystallogr D Biol Crystallogr 66: 213–221.

Barnoy, S., Baqar, S., Kaminski, R.W., Collins, T., Nemelka, K., Hale, T.L., Ranallo, R.T., and Venkatesan, M.M. (2011) Shigella sonnei vaccine candidates WRSs2 and WRSs3 are as immunogenic as WRSS1, a clinically tested vaccine candidate, in a primate model of infection. Vaccine 29: 6371–6378.

Bedford, L., Fonseka, S., Boren, T., Ranallo, R.T., Suvarnapunya, A.E., Lee, J.E., Barnoy, S., and Venkatesan, M.M. (2011) Further characterization of Shigella sonnei live vaccine candidates WRSs2 and WRSs3-plasmid composition, invasion assays and Sereny reactions. Gut Microbes 2: 244–251.

Buchrieser, C., Glaser, P., Rusniok, C., Nedjari, H., D’Hauteville, H., Kunst, F., Sansonetti, P., and Parsot, C. (2000) The virulence plasmid pWR100 and the repertoire of proteins secreted by the type III secretion apparatus of Shigella flexneri. Mol Microbiol 38: 760–771.

Caboni, M., Pedron, T., Rossi, O., Goulding, D., Pickard, D., Citiulo, F., MacLennan, C.A., Dougan, G., Thomson, N.R., Saul, A., Sansonetti, P.J., and Gerke, C. (2015) An O Antigen Capsule Modulates Bacterial Pathogenesis in Shigella sonnei. PLoS Pathog 11: e1004749.

Cintron, M., Zeng, J.M., Barth, V.C., Cruz, J.W., Husson, R.N., and Woychik, N.A. (2019) Accurate target identification for Mycobacterium tuberculosis endoribonuclease toxins requires expression in their native host. Sci Rep 9: 5949.

Datsenko, K.A., and Wanner, B.L. (2000) One-step inactivation of chromosomal genes in Escherichia coli K-12 using PCR products. Proc Natl Acad Sci U S A 97: 6640–6645.

Delageniere, S., Brenchereau, P., Launer, L., Ashton, A.W., Leal, R., Veyrier, S., Gabadinho, J., Gordon, E.J., Jones, S.D., Levik, K.E., McSweeney, S.M., Monaco, S., Nanao, M., Spruce, D., Svensson, O., Walsh, M.A., and Leonard, G.A. (2011) ISPyB: an information management system for synchrotron macromolecular crystallography. Bioinformatics 27: 3186–3192.

Dienemann, C., Bøggild, A., Winther, K.S., Gerdes, K., and Brodersen, D.E. (2011) Crystal structure of the VapBC toxin-antitoxin complex from Shigella flexneri reveals a hetero-octameric DNA-binding assembly. J Mol Biol 414: 713–722.

Ditta, G., Stanfield, S., Corbin, D., and Helinski, D.R. (1980) Broad host range DNA cloning system for gram-negative bacteria: construction of a gene bank of Rhizobium meliloti. Proc Natl Acad Sci U S A 77: 7347–7351.

Emsley, P., Lohkamp, B., Scott, W.G., and Cowtan, K. (2010) Features and development of Coot. Acta Crystallogr D Biol Crystallogr 66: 486–501.

Figurski, D.H., and Helinski, D.R. (1979) Replication of an origin-containing derivative of plasmid RK2 dependent on a plasmid function provided in trans. Proc Natl Acad Sci U S A 76: 1648–1652.

Gerke, C., Colucci, A.M., Giannelli, C., Sanzone, S., Vitali, C.G., Sollai, L., Rossi, O., Martin, L.B., Auerbach, J., Di Cioccio, V., and Saul, A. (2015) Production of a Shigella sonnei Vaccine Based on Generalized Modules for Membrane Antigens (GMMA), 1790GAHB. PLoS One 10: e0134478.

Gotfredsen, M., and Gerdes, K. (1998) The Escherichia coli relBE genes belong to a new toxin-antitoxin gene family. Mol Microbiol 29: 1065–1076.

Gronlund, H., and Gerdes, K. (1999) Toxin-antitoxin systems homologous with relBE of Escherichia coli plasmid P307 are ubiquitous in prokaryotes. J Mol Biol 285: 1401–1415.

Guzman, L.M., Belin, D., Carson, M.J., and Beckwith, J. (1995) Tight regulation, modulation, and high-level expression by vectors containing the arabinose PBAD promoter. J Bacteriol 177: 4121–4130.

Hartman, A.B., and Venkatesan, M.M. (1998) Construction of a stable attenuated Shigella sonnei DeltavirG vaccine strain, WRSS1, and protective efficacy and immunogenicity in the guinea pig keratoconjunctivitis model. Infect Immun 66: 4572–4576.

Holt, K.E., Baker, S., Weill, F.X., Holmes, E.C., Kitchen, A., Yu, J., Sangal, V., Brown, D.J., Coia, J.E., Kim, D.W., Choi, S.Y., Kim, S.H., da Silveira, W.D., Pickard, D.J., Farrar, J.J., Parkhill, J., Dougan, G., and Thomson, N.R. (2012) Shigella sonnei genome sequencing and phylogenetic analysis indicate recent global dissemination from Europe. Nat Genet 44: 1056–1059.

Jiang, Y., Yang, F., Zhang, X., Yang, J., Chen, L., Yan, Y., Nie, H., Xiong, Z., Wang, J., Dong, J., Xue, Y., Xu, X., Zhu, Y., Chen, S., and Jin, Q. (2005) The complete sequence and analysis of the large virulence plasmid pSS of Shigella sonnei. Plasmid 54: 149–159.

Kopecko, D.J., Holcombe, J., and Formal, S.B. (1979) Molecular characterization of plasmids from virulent and spontaneously occurring avirulent colonial variants of Shigella flexneri. Infect Immun 24: 580–582.

Kopecko, D.J., Washington, O., and Formal, S.B. (1980) Genetic and physical evidence for plasmid control of Shigella sonnei form I cell surface antigen. Infect Immun 29: 207–214.

Kotloff, K.L., Nataro, J.P., Blackwelder, W.C., Nasrin, D., Farag, T.H., Panchalingam, S., Wu, Y., Sow, S.O., Sur, D., Breiman, R.F., Faruque, A.S., Zaidi, A.K., Saha, D., Alonso, P.L., Tamboura, B., Sanogo, D., Onwuchekwa, U., Manna, B., Ramamurthy, T., Kanungo, S., Ochieng, J.B., Omore, R., Oundo, J.O., Hossain, A., Das, S.K., Ahmed, S., Qureshi, S., Quadri, F., Adegbola, R.A., Antonio, M., Hossain, M.J., Akinsola, A., Mandomando, I., Nhampossa, T., Acacio, S., Biswas, K., O’Reilly, C.E., Mintz, E.D., Berkeley, L.Y., Muhsen, K., Sommerfelt, H., Robins-Browne, R.M., and Levine, M.M. (2013) Burden and aetiology of diarrhoeal disease in infants and young children in developing countries (the Global Enteric Multicenter Study, GEMS): a prospective, case-control study. Lancet 382: 209–222.

Kotloff, K.L., Taylor, D.N., Sztein, M.B., Wasserman, S.S., Losonsky, G.A., Nataro, J.P., Venkatesan, M., Hartman, A., Picking, W.D., Katz, D.E., Campbell, J.D., Levine, M.M., and Hale, T.L. (2002) Phase I evaluation of delta virG Shigella sonnei live, attenuated, oral vaccine strain WRSS1 in healthy adults. Infect Immun 70: 2016–2021.

Kotloff, K.L., Winickoff, J.P., Ivanoff, B., Clemens, J.D., Swerdlow, D.L., Sansonetti, P.J., Adak, G.K., and Levine, M.M. (1999) Global burden of Shigella infections: implications for vaccine development and implementation of control strategies. Bull World Health Organ 77: 651–666.

Lan, R., and Reeves, P.R. (2002) Escherichia coli in disguise: molecular origins of Shigella. Microbes Infect 4: 1125–1132.

Levin-Reisman, I., Ronin, I., Gefen, O., Braniss, I., Shoresh, N., and Balaban, N.Q. (2017) Antibiotic tolerance facilitates the evolution of resistance. Science 355: 826–830.

Livio, S., Strockbine, N.A., Panchalingam, S., Tennant, S.M., Barry, E.M., Marohn, M.E., Antonio, M., Hossain, A., Mandomando, I., Ochieng, J.B., Oundo, J.O., Qureshi, S., Ramamurthy, T., Tamboura, B., Adegbola, R.A., Hossain, M.J., Saha, D., Sen, S., Faruque, A.S., Alonso, P.L., Breiman, R.F., Zaidi, A.K., Sur, D., Sow, S.O., Berkeley, L.Y., O’Reilly, C.E., Mintz, E.D., Biswas, K., Cohen, D., Farag, T.H., Nasrin, D., Wu, Y., Blackwelder, W.C., Kotloff, K.L., Nataro, J.P., and Levine, M.M. (2014) Shigella isolates from the global enteric multicenter study inform vaccine development. Clin Infect Dis 59: 933–941.

Mattock, E., and Blocker, A.J. (2017) How Do the Virulence Factors of Shigella Work Together to Cause Disease? Front Cell Infect Microbiol 7: 64.

Maurelli, A.T., Baudry, B., d’Hauteville, H., Hale, T.L., and Sansonetti, P.J. (1985) Cloning of plasmid DNA sequences involved in invasion of HeLa cells by Shigella flexneri. Infect Immun 49: 164–171.

Maurelli, A.T., Blackmon, B., and Curtiss, R., 3rd (1984) Loss of pigmentation in Shigella flexneri 2a is correlated with loss of virulence and virulence-associated plasmid. Infect Immun 43: 397–401.

McCoy, A.J., Grosse-Kunstleve, R.W., Adams, P.D., Winn, M.D., Storoni, L.C., and Read, R.J. (2007) Phaser crystallographic software. J Appl Crystallogr 40: 658–674.

McVicker, G., Hollingshead, S., Pilla, G., and Tang, C.M. (2019) Maintenance of the virulence plasmid in Shigella flexneri is influenced by Lon and two functional partitioning systems. Mol Microbiol 111: 1355–1366.

McVicker, G., and Tang, C.M. (2016) Deletion of toxin–antitoxin systems in the evolution of Shigella sonnei as a host-adapted pathogen. Nature Microbiology 2: 16204.

Ogura, T., and Hiraga, S. (1983) Mini-F plasmid genes that couple host cell division to plasmid proliferation. Proc Natl Acad Sci U S A 80: 4784–4788.

Overgaard, M., Borch, J., Jorgensen, M.G., and Gerdes, K. (2008) Messenger RNA interferase RelE controls relBE transcription by conditional cooperativity. Mol Microbiol 69: 841–857.

Payne, S.M., and Finkelstein, R.A. (1977) Detection and differentiation of iron-responsive avirulent mutants on Congo red agar. Infect Immun 18: 94–98.

Pilla, G., McVicker, G., and Tang, C.M. (2017) Genetic plasticity of the Shigella virulence plasmid is mediated by intra- and inter-molecular events between insertion sequences. PLoS Genet 13: e1007014.

Pullinger, G.D., and Lax, A.J. (1992) A Salmonella dublin virulence plasmid locus that affects bacterial growth under nutrient-limited conditions. Mol Microbiol 6: 1631–1643.

Radnedge, L., Davis, M.A., Youngren, B., and Austin, S.J. (1997) Plasmid maintenance functions of the large virulence plasmid of Shigella flexneri. J Bacteriol 179: 3670–3675.

Robson, J., McKenzie, J.L., Cursons, R., Cook, G.M., and Arcus, V.L. (2009) The vapBC operon from Mycobacterium smegmatis is an autoregulated toxin-antitoxin module that controls growth via inhibition of translation. J Mol Biol 390: 353–367.

Sansonetti, P., David, M., and Toucas, M. (1980) [Correlation between the loss of plasmid DNA and the transition from virulent phase I to avirulent phase II in Shigella sonnei]. C R Seances Acad Sci D 290: 879–882.

Sansonetti, P.J., Kopecko, D.J., and Formal, S.B. (1981) Shigella sonnei plasmids: evidence that a large plasmid is necessary for virulence. Infect Immun 34: 75–83.

Sansonetti, P.J., Ryter, A., Clerc, P., Maurelli, A.T., and Mounier, J. (1986) Multiplication of *Shigella flexneri* within HeLa cells: lysis of the phagocytic vacuole and plasmid-mediated contact hemolysis. Infect Immun 51: 461–469.

Sasakawa, C., Kamata, K., Sakai, T., Makino, S., Yamada, M., Okada, N., and Yoshikawa, M. (1988) Virulence-associated genetic regions comprising 31 kilobases of the 230-kilobase plasmid in Shigella flexneri 2a. J Bacteriol 170: 2480–2484.

Sasakawa, C., Kamata, K., Sakai, T., Murayama, S.Y., Makino, S., and Yoshikawa, M. (1986) Molecular alteration of the 140-megadalton plasmid associated with loss of virulence and Congo red binding activity in Shigella flexneri. Infect Immun 51: 470–475.

Sayeed, S., Brendler, T., Davis, M., Reaves, L., and Austin, S. (2005) Surprising dependence on postsegregational killing of host cells for maintenance of the large virulence plasmid of Shigella flexneri. J Bacteriol 187: 2768–2773.

Sayeed, S., Reaves, L., Radnedge, L., and Austin, S. (2000) The stability region of the large virulence plasmid of Shigella flexneri encodes an efficient postsegregational killing system. J Bacteriol 182: 2416–2421.

Schroeder, G.N., and Hilbi, H. (2008) Molecular pathogenesis of *Shigella* spp.: controlling host cell signaling, invasion, and death by type III secretion. Clin Microbiol Rev 21: 134–156.

Schuch, R., and Maurelli, A.T. (1997) Virulence plasmid instability in Shigella flexneri 2a is induced by virulence gene expression. Infect Immun 65: 3686–3692.

Tsai, C.N., and Coombes, B.K. (2019) The Role of the Host in Driving Phenotypic Heterogeneity in Salmonella. Trends Microbiol 27: 508–523.

Venkatesan, M.M., Goldberg, M.B., Rose, D.J., Grotbeck, E.J., Burland, V., and Blattner, F.R. (2001) Complete DNA sequence and analysis of the large virulence plasmid of Shigella flexneri. Infect Immun 69: 3271–3285.

Winn, M.D., Ballard, C.C., Cowtan, K.D., Dodson, E.J., Emsley, P., Evans, P.R., Keegan, R.M., Krissinel, E.B., Leslie, A.G., McCoy, A., McNicholas, S.J., Murshudov, G.N., Pannu, N.S., Potterton, E.A., Powell, H.R., Read, R.J., Vagin, A., and Wilson, K.S. (2011) Overview of the CCP4 suite and current developments. Acta Crystallogr D Biol Crystallogr 67: 235–242.

Winter, G., Waterman, D.G., Parkhurst, J.M., Brewster, A.S., Gildea, R.J., Gerstel, M., Fuentes-Montero, L., Vollmar, M., Michels-Clark, T., Young, I.D., Sauter, N.K., and Evans, G. (2018) DIALS: implementation and evaluation of a new integration package. Acta Crystallogr D Struct Biol 74: 85–97.

Winther, K.S., Brodersen, D.E., Brown, A.K., and Gerdes, K. (2013) VapC20 of Mycobacterium tuberculosis cleaves the sarcin-ricin loop of 23S rRNA. Nat Commun 4: 2796.

Winther, K.S., and Gerdes, K. (2011) Enteric virulence associated protein VapC inhibits translation by cleavage of initiator tRNA. Proc Natl Acad Sci U S A 108: 7403–7407.

Winther, K.S., and Gerdes, K. (2012) Regulation of enteric vapBC transcription: induction by VapC toxin dimer-breaking. Nucleic Acids Res 40: 4347–4357.

Xu, D.Q., Cisar, J.O., Ambulos, N., Jr., Burr, D.H., and Kopecko, D.J. (2002) Molecular cloning and characterization of genes for Shigella sonnei form I O polysaccharide: proposed biosynthetic pathway and stable expression in a live salmonella vaccine vector. Infect Immun 70: 4414–4423.

Yanisch-Perron, C., Vieira, J., and Messing, J. (1985) Improved M13 phage cloning vectors and host strains: nucleotide sequences of the M13mp18 and pUC19 vectors. Gene 33: 103–119.

Zychlinsky, A., Prevost, M.C., and Sansonetti, P.J. (1992) Shigella flexneri induces apoptosis in infected macrophages. Nature 358: 167–169.

