## Supplementary Tables and Figures for "The role of toxin:antitoxin systems and insertion sequences in the loss of virulence in *Shigella sonnei*"

**Supplementary Table 1: The presence of *relBE* on pINV$^{Ssonn}$**

| Sequence ID Holt *et al.* | Accession number | *relBE* | ori | *vapBC* |
|---|---|---|---|---|
| Sh74369 | SRA: ERX2006371 | present | present | present |
| CS1 | NZ_CXFA00000000 | present | - | present |
| 20071599 | NZ_CXET00000000 | present | present | present |
| Sh9810267 | NZ_CXDR00000000 | present | present | present |
| ShIB716 | SRA:ERX2005632 | present | present | present |
| ShIB717 | NZ_CXBM00000000 | present | present | present |
| ShIB748 | NZ_CXBQ00000000 | present | present | present |
| ShIB697 | NZ_CXBP00000000 | present | present | present |
| Sh2073 | SRA: ERX2005582 | present | present | present |
| CS8 | NZ_CXEZ00000000 | - | - | - |
| ShIB3277 | SRA: ERX2005656 | present | present | present |
| ShIB3580 | NZ_CXBZ00000000 | present | present | present |
| ShIB3599 | NZ_CXCP00000000 | present | present | present |
| ShIB3488 | NZ_CXBW00000000 | present | present | present |
| ShIB3507 | NZ_CXBT00000000 | present | present | present |
| CS7 | NZ_CXEY00000000 | present | present | present |
| 20010007 | NZ_CXEX00000000 | present | present | present |
| 20062313 | NZ_CXEW00000000 | present | present | present |
| ShIB3300 | NZ_CXCL00000000 | present | present | present |
| CS20 | NZ_CXEE00000000 | present | present | present |
| CS14 | NZ_CXEI00000000 | present | present | present |
| 19904011 | NZ_CXES00000000 | present | present | present |
| Sh970044 | SRA: ERS009863 | present | present | present |
| 20011685 | NZ_CXER00000000 | present | present | present |
| 20031275 | NZ_CXEM00000000 | present | present | present |
| Sh60108 | NZ_CXAS00000000 | present | present | present |
| 20062087 | NZ_CXEU00000000 | present | present | present |
| ShIB691 | NZ_CXAZ00000000 | present | present | present |
| 20040924 | NZ_CXEV00000000 | present | present | present |
| ShIB3374 | SRA: ERS009826 | present | present | present |
| 20021122 | NZ_CXEG00000000 | present | present | present |
| ShIB2013 | NZ_CXBN00000000 | present | present | present |
| ShIB2012 | SRA:ERX2005602 | present | present | present |
| ShIB1976 | SRA: ERS009804 | present | present | present |
| ShIB1980 | NZ_CXBI00000000 | present | present | present |
| ShIB1985 | NZ_CXCG00000000 | present | present | presenrt |
| ShIB2015 | NZ_CXCF00000000 | present | present | present |
| ShIB2008 | NZ_CXBV00000000 | present | present | present |
| ShIB2000 | NZ_CXBY00000000 | present | present | present |
| ShIB1970 | NZ_CXBL00000000 | present | present | present |

| | | | | |
|---|---|---|---|---|
| ShIB2004 | NZ_CXCD00000000 | present | present | present |
| ShIB2024 | NZ_CXCA00000000 | present | present | present |
| ShIB2018 | NZ_CXBU00000000 | present | present | present |
| ShIB1993 | NZ_CXBO00000000 | present | present | present |
| ShIB1990 | NZ_CXCK00000000 | present | present | present |
| ShIB1987 | NZ_CXBR00000000 | present | present | present |

**Supplementary Table 2: Polymorphisms in VapB from *S. flexneri* and *S. sonnei.*** The frequency of each polymorphism is shown as a percentage of the total.

| | Amino acid sequence | Percentage of VapB sequences with given amino acid sequence |
|---|---|---|
| ***S. sonnei* VapB** | | Total=336 sequences |
| | METTVFLSNRSQAVRLPKAVALPENVKRVEVIAVGRTRIITPAGETWDEWFDGHSVSADFMDNREQPGMQERESF* | 87.61329 Including *S. sonnei* 53G |
| | METTVFLSNRSQAVRLPKAVALPENVKRVEVIAIGRTRLITPAGETWDEWFDGHSVSADFMDNREQPGMQERESF* | 0.302115 |
| | METTVFLSNRSQAVRLPKAVALPENVKRVEVIAVGRTRIITPAGETWDEWFDGHSVSTDFMDNREQPGMQERESF* | 0.302115 |
| | METTVFLSNRSQAVRLPKAVALPENVKRVEVIAVGRTRIITPAGETWDEWFDGNSVSADFMDNREQPGMQERESF* | 0.60423 |
| | METTVFLSNRSLAVRLPKAVALPENVKRVEVIAVGRTRIITPAGETWDEWFDGHSVSADFMDNREQPGMQERESF* | 0.302115 |
| | METTVFFSNRSQAVRLPKAVALPENVKRVEVIAVGRTRIITPAGETWDEWFDGHNVSADFMDIRDQPAMQERESF* | 2.114804 |
| | METTVFLSNRSQAVRLPKAVALPEDVKKVEIIAIGRTRIITPAGESWDSWFDGENVSADFMDIRDQPAMQERESF* | 8.761329 |
| ***S. flexneri* VapB** | | Total=310 sequences |
| | METTVFLSNRSQAVRLPKAVALPENVKRVEVIAVGRTRIITPAGETWDEWFDGHSVSTDFMDNREQPGMQERESF* | 95.48387 Including *S. flexneri* M90T |
| | METTVFLSNRSQAVRLPKAVALPENVKRVEVIAVGRTRIITPAGETWDEWFDGHSVSADFMDNREQPGMQERESF* | 4.516129 |

**Supplementary Table 3: Polymorphisms in VapC from *S. flexneri* and *S. sonnei*.** The frequency of each polymorphism is shown as a percentage of the total.

| | Amino acid sequence | Percentage of VapC sequences with given amino acid sequence |
|---|---|---|
| ***S. sonnei* VapC** | | Total= 320 sequences |
| | MLKFMLDTNICIFTIKNKPASVRERFNLNQGRMCISSVTLMELIYGAEKSQMPERNLAVIEGFVSRIDVLDYDAAAATHTGQIRAELARQGRPVGPFDQMIAGHARSRGLIIVTNNTREFERVGGLRTEDWS* | 82.8125 Including *S. sonnei* 53G |
| | MLKFMLDTNICIFTIKNKPASIRERFNLNQGRMCISSVTLMELIYGAEKSQMPERNLAVIEGFVSRIDVLDYDAAAATHTGQIRAELARQGRPVGPFDQMIAGHARSRGLIIVTNNTREFERVGGLRTEDWS* | 2.5 |
| | MLKFMLDTNICIFTIKNKPASVRERFNLNQGRMCISSVTLMELIYGAEKSQMPERNLAVIEGFASRIDVLDYDAAAATHTGQIRAELARQGRPVGPFDQMIAGHARSRGLIIVTNNTREFERVGGLRTEDWS* | 0.3125 |
| | MLKFMLDTNICIFTIKNNPASVRERFNLNQGRMCISSVTLMELIYGAEKSQMPERNLAVIEGFVSRIDVLDYDAAAATHTGQIRAELARQGRPVGPFDQMIAGHARSRGLIIVTNNTREFERVGGLRTEDWS* | 0.9375 |
| | MLKFMLDTNICIFTIKNKPASVRERFNLNQGRMCISSVTLMELIYGAEKSQMPERNLAVIEGFVSRIDVLDYDAAAATHTGQIRAELARQGRPVGPFDQMIAGHARSRGLIIVTNSTREFERVGGLRTEDWS* | 0.3125 |
| | MLKFMLDTNICIFTIKNKPASVRERFNLNQGRMCISSVTLMELIYGAEKSQMPERNLAVIEGFVSRIDVLDYDAAAAIHTGQIRAELARQGRPVGPFDQMIAGHARSRGLIIVTNNTREFERVGGLRTEDWS* | 0.625 |
| | MLKFMLDTNICIFTIKNKPASVRERFNLNQGRMCISSVTLMELIYGAEKSQMPERNLAVIEGFVSRIDVLDYDAAAATHSGQIRAELARQGRPVGPFDQMIAGHARSRGLIIVTNNTREFERVDGLRIEDWS* | 1.875 |
| | MLKFMLDTNICIFTIKNKPASVRERFNLNQGRMCISSVTLMEVIYGAEKSQMPERNLAVIEGFVSRIDVLDYDAAAATHTGQIRAELALQGRPVGPFDQMIAGHARSRGLIIVTNNTREFERVGGLRIEDWS* | 0.625 |
| | MLKFMLDTNICIFTIKNKPASVRERFNLNQGRMCISSVTLMELIYGAEKSQMPERNLAVIEGFVSRIDVLDYDAAAAIHTGQIRAELARQGRPVGPFDQMIAGHARCRGLIIVTNNTREFERVGGLRIEDWS* | 0.3125 |
| | MLKFMLDTNICIFTMKNKPASVRERFNLNQGRMCISSVTLMELIYGAEKSQMPERNLAVIEGFVSRIDVLDYDAAAATHSGQIRAELARQGRPVGPFDQMIAGHARSRGLIIVTNNTREFERVDGLRIEDWS* | 0.3125 |
| | MLKFMLDTNICIFTIKNKPVHVRERFNLNSGRMCISSVTLMELIYGAEKSQMPERNLAVIEGFVSRLEVLDYDSAAATHTGQIRAELARQGRPVGPFDQMIAGHARSRGLIVVTNNTREFERVAGIRLEDWS* | 3.125 |
| | MLKFMLDTNICIFTIKNKPVHVRERFNLNSGRMCISSVTLMELIYGAEKSQMPERNLAVIEGFVSRLVVLDYDSAAATHTGQIRAELARQGRPVGPFDQMIAGHARSRGLIVVTNNTREFERVAGIRLEDWS* | 5.625 |
| | MLKFMLDTNICIFTIKNKPVHVRERFNLNSGRMCISSVTLVELIYGAEKSQMPERNLAVIEGFVSRLVVLDYDSAAATHTGQIRAELARQGRPVGPFDQMIAGHARSRGLIVVTNNTREFERVAGIRLEDWS* | 0.625 |
| ***S. flexneri* VapC** | | Total= 519 sequences |
| | MLKFMLDTNICIFTIKNKPASVRERFNLNQGKMCISSVTLMELIYGAEKSQMPERNLAVIEGFVSRIDVLDYDAAAATHTGQIRAELARQGRPVGPFDQMIAGHARSRGLIIVTNNTREFERVGGLRTEDWS* | 78.22736 Including *S. flexneri* M90T |
| | MLKFMLDTNICIFTIKNKPASVRERFNLNQGRMCISSVTLMELIYGAEKSQMPERNLAVIEGFVSRIDVLDYDAAAATHTGQIRAELARQGRPVGPFDQMIAGHARSRGLIIVTNNTREFERVGGLRTEDWS* | 6.358382 |
| | MLKFMLDTNICIFTIKNKPASVRERFNLNQGKMCISSVTLMKLIYGAEKSQMPERNLAVIEGFVSRIDVLDYDAAAATHTGQIRAELARQGRPVGPFDQMIAGHARSRGLIIVTNNTREFERVGGLRTEDWS* | 0.385356 |
| | MLKFILDTNICIFTIKNKPASVRERFNLNQGKMCISSVTLMELIYGAEKSQMPERNLAVIEGFVSRIDVLDYDAAAATHTGQIRAELARQGRPVGPFDQMIAGHARSRGLIIVTNNTREFERVGGLRTEDWS* | 8.863198 |
| | MLKFMLDTNICIFTIKNKPASVRERFNLNQGKMCISLVTLMELIYGAEKSQMPERNLAVIEGFVSRIDVLDYDAAAATHTGQIRAELARQGRPVGPFDQMIAGHARSRGLIIVTNNTREFERVGGLRTEDWS* | 0.192678 |
| | MLKFMLNTNICIFTIKNKPASVRERFNLNQGKMCISSVTLMELIYGAEKSQMPERNLAVIEGFVSRIDVLDYDAAAATHTGQIRAELARQGRPVGPFDQMIAGHARSRGLIIVTNNTREFERVGGLRTEDWS* | 2.312139 |
| | MLKFMLDTNICIFTIKNKPASVRERFNLNQGKMCISSVTLMELIYGAEKSQMPERNLAVIEGFVSRIDVLDYDAAAATHTGQIRAELARQGRPVGPFNQMIAGHARSRGLIIVTNNTREFERVGGLRTEDWS* | 1.926782 |
| | MLKFMLDTNICIFTIKNKPASVRERFNLNQGRMCISSVTLMELIYGAEKSQMSERNLAVIEGFVSRIDVLDYDAAAATHTGQIRAELARQGRPVGPFDQMIAGHARSRGLIIVTNNTREFERVGGLRTEDWS* | 0.578035 |
| | MLKFMLDTNICIFTIKNKPASVRERFNLNQGRMCISSVTLMELIYGAEKSQMPERNLAVIEGFVSRIDVLDYDAAAATHSGQIRAELARQGRPVGPFDQMIAGHARSRGLIIVTNNTREFERVDGLRIEDWS* | 0.192678 |
| | MLKFMLDTNICIFTIKNKPVHVRERFNLNSGRMCISSVTLMELIYGAEKSQMPERNLAVIEGFVSRLVVLDYDSAAATHTGQIRAELARQGRPVGPFDQMIAGHARSRGLIVVTNNTREFERVAGIRLEDWS* | 0.963391 |

**Supplementary Table 4: Bacterial strains used in this study**

| Strain name | Relevant genotype/description | Reference |
|---|---|---|
| *S. flexneri* M90T | Wild-type *S. flexneri* serotype 5a | Zychlinsky *et al.* 1992 |
| BS176 | *S. flexneri* M90T lacking pINV | Zychlinsky *et al.* 1992 |
| *S. sonnei* 53G | Wild-type *S. sonnei* | Kopecko *et al.* 1980 |
| *S. flexneri vapBC-cat* | *S. flexneri* with *cat* downstream of *vapBC* | Pilla *et al*., 2017 |
| *S. sonnei vapBC-cat* | *S. sonnei* with *cat* downstream of *vapBC* | This study |
| *S. sonnei* $pINV^-$ | *S. sonnei* lacking pINV | McVicker & Tang, 2016 |
| *S. sonnei* Δ*relBE* | *S. sonnei* Δ*relBE mxiH::sacB-neo* | This study |
| *S. sonnei* $ccdAB^+/gmvAT^+$ | *S. sonnei* containing *ccdAB-cat gmvAT* | McVicker & Tang, 2016 |
| *S. sonnei* $VapBC^{Sflex}$ | *S. sonnei* containing $vapBC^{Sflex}$ with *cat* downstream | This study |
| *S. sonnei* $VapB^{Ssonn}C^{Sflex}$ | *S. sonnei* containing $vapB^{Ssonn}C^{Sflex}$ with *cat* downstream | This study |
| *S. sonnei* $VapB^{Sflex}C^{Ssonn}$ | *S. sonnei* containing $vapB^{Sflex}C^{Ssonn}$ with *cat* downstream | This study |
| BS176 $pINV^{Ssonn}$ $VapBC^{Ssonn}$ | BS176 containing $pINV^{Ssonn}$ $VapBC^{Ssonn}$ with *cat* downstream | This study |
| BS176 $pINV^{Ssonn}$ $VapBC^{Sflex}$ | BS176 containing $pINV^{Ssonn}$ $VapBC^{Sflex}$ with *cat* downstream | This study |
| *E. coli* DH5α | *fhuA*2 Δ(*argF-lacZ*) *U169 phoA glnV44* Φ*80* Δ(*lacZ*)*M15 gyrA96 recA1 relA1 endA1 thi-1 hsdR17* | Hanahan 1983 |
| *E. coli* MG1655 | Wild-type *E. coli* K-12 | Guyer *et al.* 1981 |
| *E. coli* C41(DE3) | *ompT hsdSB* (*rB- mB-*) *gal dcm* (DE3) | Miroux & Walker, 1996 |

**Supplementary Table 5: Plasmids used in this study**

| Plasmid | Relevant genotype or description | Antibiotic resistance | Reference |
|---|---|---|---|
| pUC19 | Cloning vector | $Amp^r$ | Yanisch-Perron *et al.* 1985 |
| pUC19-*relBE*::*cat* | Cloning vector to introduce *relBE* with a downstream *cat* cassette into *S. sonnei* | $Amp^r$ | This study |
| pUC19-*vapBC::cat* | Cloning vector to introduce *vapBC* with a downstream *cat* cassette into *S. sonnei* | $Amp^r$ | This study |
| pUC19-*vapBC*$^{Sflex}$ | Cloning vector to introduce *vapBC* from *S. flexneri* with a downstream *cat* cassette into *S. sonnei* | $Amp^r$ | This study |
| pUC19-*vapB*$^{Ssonn}$*C*$^{Sflex}$ | to introduce *vapC* from *S. flexneri* into *S. sonnei vapBC-cat* | $Amp^r$ | This study |
| pUC19-*vapB*$^{Sflex}$*C*$^{Ssonn}$ | Cloning vector to introduce *vapB* from *S. flexneri* into *S. sonnei vapBC-cat* | $Amp^r$ | This study |
| pKD46 | Plasmid expressing the λ red recombinase | $Amp^r$ | Datsenko & Wanner, 2000 |
| pCP20 | Temperature sensitive plasmid encoding FLP recombinase | $Amp^r$ | Datsenko & Wanner, 2000 |
| pKD3 | Plasmid with FRT-sites flanking a *cat* cassette | $Kan^r$ | Datsenko & Wanner, 2000 |
| pSTAB | 6.3 kb plasmid containing the pINV ori and *sacB-neo* | $Kan^r$ | McVicker *et al.* 2019 |
| pSTAB::VapBC$^{Sflex}$ | pSTAB with *S. flexneri vapBC* | $Kan^r$ | This study |
| pSTAB::VapBC$^{Ssonn}$ | pSTAB with *S. sonnei vapBC* | $Kan^r$ | This study |
| pSTAB::VapB$^{Ssonn}$C$^{Sflex}$ | pSTAB with *vapB* from *S. sonnei* and *vapC* from *S. flexneri* | $Kan^r$ | This study |
| pSTAB::VapB$^{Sflex}$C$^{Ssonn}$ | pSTAB with *vapB* from *S. flexneri* and *vapC* from *S.sonnei* | $Kan^r$ | This study |
| pBAD33 | Empty vector | $Cam^r$ | Guzman *et al.* 1995 |
| pBAD33::RelE | pBAD33 with *relE* under an arabinose-inducible promoter | $Cam^r$ | This study |
| pBAD33::VapC$^{Sflex}$ | pBAD33 with *S. flexneri vapC* under an arabinose-inducible promoter | $Cam^r$ | McVicker & Tang, 2016 |
| pBAD33::VapC$^{Sflex}$ D7A | pBAD33 with *S. flexneri vapC*$^{D7A}$ under an arabinose-inducible promoter | $Cam^r$ | McVicker & Tang, 2016 |
| pBAD33::VapC$^{Ssonn}$ | pBAD33 with *S. sonnei vapC* under an arabinose-inducible promoter | $Cam^r$ | This study |
| pGM101 | Vector compatible with pBAD33 | $Amp^r$ | McVicker & Tang, 2016 |
| pGM101::RelB | pGM101 with *relB* under its native promoter | | This study |
| pGM101::VapB$^{Sflex}$ | pGM101 with *vapB* from *S. flexneri* under its native promoter | $Amp^r$ | McVicker & Tang, 2016 |
| pGM101::VapB$^{Ssonn}$ | pGM101 with *vapC* from *S. sonnei* under its native promoter | $Amp^r$ | This study |
| pINV$^{Ssonn}$ VapBC$^{Ssonn}$ | *S. sonnei* pINV with *vapBC*$^{Ssonn}$*-cat* | $Cam^r$ | This study |
| pINV$^{Ssonn}$ VapBC$^{Sflex}$ | *S. sonnei* pINV with *vapBC*$^{Sflex}$*-cat* | $Cam^r$ | This study |
| pET28a-VapBC$^{Ssonn}$ | pET28a with N-terminal His tagged VapC and untagged VapB from *S. sonnei* | $Kan^r$ | This study |

**Supplementary Table 6: Primers used in this study**

| Primer | Sequence (5' - 3') | Purpose |
|---|---|---|
| GM371 | tgggctagcgaattcgagctcatgatgtggacctggac | Construction of pBAD33-*relE* |
| GM372 | ctcatccgccaaaacagccaagcttctatctcattctctcactgg | Construction of pBAD33-*relE* |
| GM373 | cgcgaaggcgaagcggcatgcgctgaaaacgggtatggc | Construction of pGM101-*relB* |
| GM374 | ctcatccgccaaaacagccaagcttcataaataactgtccaggtcc | Construction of pGM101-*relB* |
| GM399 | aaacgacggccagtgaattcgagctcaagtgacctcctcagaataatc | Construction of pUC19-*relBE::cat* |
| GM400 | cttcgaagcagctccagcctacacattaacaagtctttattactcacac | Construction of pUC19-*relBE::cat* |
| GM401 | catatggaccatggctaattcccattaacatataaataacctttcccg | Construction of pUC19-*relBE::cat* |
| GM402 | aacagctatgaccatgattacgccaagcttgtacatgtgtctgtaac | Construction of pUC19-*relBE::cat* |
| GP007 | acggccagtgaattcgagctcgtgaagcgggtccgggtg | Construction of *S. sonnei vapBC-cat, S. sonnei* vapBC$^{Sflex}$, *S. sonnei* vapB$^{Ssonn}$C$^{Sflex}$, *S. sonnei* vapB$^{Sflex}$C$^{Ssonn}$ |
| GP010 | ccatggctaattcccattcagctccagtcttcagttc | Construction of *S. sonnei vapBC-cat, S. sonnei* VapBC$^{Sflex}$, *S. sonnei* vapB$^{Ssonn}$C$^{Sflex}$, *S. sonnei* vapB$^{Sflex}$C$^{Ssonn}$ |
| GP011 | agcctacacacctgttcatcagaaatcatctc | Construction of *S. sonnei vapBC-cat, S. sonnei* vapBC$^{Sflex}$, *S. sonnei* vapB$^{Ssonn}$C$^{Sflex}$, *S. sonnei* vapB$^{Sflex}$C$^{Ssonn}$ |
| GP141 | catgattacgccaagctgtaatcgtcgtgtgtgg | Construction of *S. sonnei vapBC-cat, S. sonnei* vapBC$^{Sflex}$, *S. sonnei* vapB$^{Ssonn}$C$^{Sflex}$, *S. sonnei* vapB$^{Sflex}$C$^{Ssonn}$ |
| GM174 | tgtgtaggctggagctgctt | PCR FRT flanked *cat* cassette from pKD3 |
| GM175 | atgggaattagccatggtcc | PCR FRT flanked *cat* cassette from pKD3 |
| GM393 | ccatatggctagcatgagcggatccgtaattatccttgtcggtataataattacc | Construction of pSTAB-VapBC plasmids |
| GM394 | ggtgctcgagtgcggccgcaagcttgaacaggtcagctccag | Construction of pSTAB-VapBC plasmids |
| GM105 | aggcgaagcggcatgcgatgcctgttactgcca | pGM101-*vapB*$^{Ssonn}$, pGM101-*vapB*$^{Sflex}$ |
| GM106 | ccgccaaaacagccatcagaatgactccctttcc | pGM101-*vapB*$^{Ssonn}$, pGM101-*vapB*$^{Sflex}$ |
| GM122 | ctagcgaattcgagctcatgcaggaaagggagtc | pBAD33-*vapC*$^{Ssonn}$, pBAD33-*vapC*$^{Sflex}$ |
| GM123 | ccgccaaaacagccaagcttcagctccagtcttcagttc | pBAD33-*vapC*$^{Ssonn}$, pBAD33-*vapC*$^{Sflex}$ |
| GP99 | cctatcgatggacatgaatgcagg | Check of IS*21*-mediated deletion of T3SS-related genes |
| GP100 | tcgcctgtttcaccgtatcttc | Check of IS*21*-mediated deletion T3SS-related genes |
| GP145 | ggagcggtatcaggcaaacg | Check of IS*1294*-mediated deletion T3SS-related genes |
| GP146 | gctgacagtggtaatacgttg | Check of IS*1294*-mediated deletion T3SS-related genes |
| GP147 | ccgaccatatccgccattg | Check of IS*1*-mediated deletion T3SS-related genes |

| | | |
|---|---|---|
| GP148 | cgtttgcctgataccgctcc | Check of IS*1*-mediated deletion T3SS-related genes |
| *hns*_F | ttgcaaaggcgttgaatta | Multiplex PCR |
| *hns*_R | ttgcaaaggcgttgaatta | Multiplex PCR |
| *virB*_F | acatcagagctccacaagaa | Multiplex PCR |
| *virB*_R | agacgatagatggcgagaaa | Multiplex PCR |
| *virF*_F | cttagcttgttgcacagaga | Multiplex PCR |
| *virF*_R | aagatgggcttgatattccg | Multiplex PCR |
| *ori*_F | gtgacctcctcagaataatcc | Multiplex PCR |
| *ori*_R | aaaagatacattgcaccctgt | Multiplex PCR |
| SH10 | ctggtgccgcgcggcagccatatgctgaagtttatcgtcg | Construction of pET28a-$vapCB^{Ssonn}$ |
| SH11 | ccattatttgcctccttatcagctccagtcttcagttc | Construction of pET28a-$vapCB^{Ssonn}$ |
| SH12 | ctggagctgataaggaggcaaataatggaaaccaccgtatttc | Construction of pET28a-$vapCB^{Ssonn}$ |
| SH13 | agtggtggtggtggtggtgctcgagtcagaatgactccctttc | Construction of pET28a-$vapCB^{Ssonn}$ |

**References**


Datsenko KA, Wanner BL. One-step inactivation of chromosomal genes in *Escherichia coli* K-12 using PCR products. Proc Natl Acad Sci U S A. 2000 97:6640-5.

Guyer MS, Reed RE, Steitz T, Low KB. Identification of a sex-factor-affinity site in *E. coli* as gamma delta. Cold Spr. Harb. Symp. Quant. Biol. 1981 45:135-140

Guzman LM, Belin D, Carson MJ, Beckwith J. Tight regulation, modulation, and high-level expression by vectors containing the arabinose PBAD promoter. J Bacteriol. 1995 177:4121-30

Hanahan D. Studies on transformation with *Escherichia coli*. J Mol Biol. 1083 166:557-80.

Holt, K.E., Baker, S., Weill, F.X., Holmes, E.C., Kitchen, A., Yu, J., Sangal, V., Brown, D.J., Coia, J.E., Kim, D.W., Choi, S.Y., Kim, S.H., da Silveira, W.D., Pickard, D.J., Farrar, J.J., Parkhill, J., Dougan, G., and Thomson, N.R. (2012) *Shigella sonnei* genome sequencing and phylogenetic analysis indicate recent global dissemination from Europe. Nat Genet 44: 1056-1059.

Kopecko DJ, Washington O, Formal SB. Genetic and physical evidence for plasmid control of *Shigella sonnei* form I cell surface antigen. Infect Immun. 1980 29:207-14.

McVicker G, Tang CM. Deletion of toxin–antitoxin systems in the evolution of *Shigella sonnei* as a host-adapted pathogen. Nature Microbiology. 2016 2:16204.

McVicker G, Hollingshead S, Pilla G, Tang CM. Maintenance of the virulence plasmid in *Shigella flexneri* is influenced by Lon and two functional partitioning systems. Mol Microbiol. 2019 111:1355-66.

Miroux B, Walker JE. Over-production of proteins in *Escherichia coli*: mutant hosts that allow synthesis of some membrane proteins and globular proteins at high levels. J Mol Biol. 1996 260(3):289-298.

Sansonetti PJ, Kopecko DJ, Formal SB. *Shigella sonnei* plasmids: evidence that a large plasmid is necessary for virulence. Infect Immun. 1981 34:75-83.

Yanisch-Perron C, Vieira J, Messing J. Improved M13 phage cloning vectors and host strains: nucleotide sequences of the M13mp18 and pUC19 vectors. Gene. 1985 33:103-19.

Zychlinsky A, Prevost MC, Sansonetti PJ. *Shigella flexneri* induces apoptosis in infected macrophages. Nature. 1992 358:167-9.

```
                                                                    -35                    -10
                                                                 putative relO2                  putative relO1                  M
relBE promoter p307          -GGTGATTCTGTCCATCTGTACAAAAAACAATAAAAGACTTGTTAACAGGTCATGTAAGGAGTATCTTTGAGACTGGTTAAACAGTCTTGAAAGGTGGCCTATG   103
relBE promoter pINV^Ssonn    AGCATTTAATGCATTTGCAAAAGTGTGAGTAATAAAGACTTGTTAAGTGGTCGTTTCAGGGATATCGTTAAGACTTGTTAAACAGTCTCAAAGGTGCGTTATG
                             103
                              *    *  **    *    *      *  *  *************  **** * * ***  **** ** ***** ************  ** *   *
```

**Supplementary Figure 1 Alignment of RelBE promoter**

Alignment of the promoter region of *relBE* from *E. coli* p307 (M26308) and from *S. sonnei* pINV (NC_016833) by CLUSTAL O (1.2.4). Asterisks indicate identical nucleotides. The predicted -10 and -35 are highlighted in bold. The putative *relO* sites are indicated highlighted. The first codon of RelB is indicated using a M.

**A**

```
                                                            -35                   -10
                                                         vapO2                                  vapO1                          M
vapBC promoter pINVSflex   CCTGCGCGATACTCATCATAAACGTATATCCCTTTGACATATCCCGGTATCAATCCCACAATAGATATACACAAGACATATCCACATAAGGAGGCAAATAATG  103
vapBC promoter pINVSsonn   CCTGCGCGATACTCATCATAAACGTATATCCCTTTGACATATCCCGGTATCAATCCCACAATAGATATACACAAGACATATCCACATAAGGAGGCAAATAATG  103
                           *******************************************************************************************************
```

**B**

```
VapBSflex   METTVFLSNRSQAVRLPKAVALPENVKRVEVIAVGRTRIITPAGETWDEWFDGHSVSTDFMDNREQPGMQERESF*   75
VapBSsonn   METTVFLSNRSQAVRLPKAVALPENVKRVEVIAVGRTRIITPAGETWDEWFDGHSVSADFMDNREQPGMQERESF*   75
            *********************************************************:*****************
```

**C**

```
VapCSflex   MLKFMLDTNICIFTIKNKPASVRERFNLNQGKMCISSVTLMELIYGAEKSQMPERNLAVIEGFVSRIDVLDYDAAAATHTGQIRAELARQGRPVGPFDQMIAGHARSRGLIIVTNNTREFERVGGLRTEDWS*  132
VapCSsonn   MLKFMLDTNICIFTIKNKPASVRERFNLNQGRMCISSVTLMELIYGAEKSQMPERNLAVIEGFVSRIDVLDYDAAAATHTGQIRAELARQGRPVGPFDQMIAGHARSRGLIIVTNNTREFERVGGLRTEDWS*  132
            *******************************:****************************************************************************************************
```

**Supplementary Figure 2 Polymorphisms in VapBC.**

Alignment of the *vapBC* promoter (A) VapB (B) and VapC (C) from *S. flexneri* 5a M90T pINV (AL391753) and *S. sonnei* 53G pINV (NC_016833) by CLUSTAL O (1.2.4). Polymorphic residues are shown in red. Asterisks indicate identical nucleotides/amino acids. The predicted -10 and -35 are highlighted in bold. The putative *vapO* sites are indicated.
